# Preferential formation of NUP98::KDM5A condensates at specific H3K4me3-rich loci drives leukemogenic gene expression

**DOI:** 10.64898/2026.03.16.712262

**Authors:** Augusto Berrocal, Jonathan E Sandoval, Neha Khetan, Alexa Ma, Tianhong Wang, Camille Moore, Geeta J Narlikar, Hao Li, Danica Galonić Fujimori, Bo Huang

## Abstract

Chromosomal translocations involving NUP98 generate fusion proteins that alter gene expression programs, yet the fundamental principles governing their gene targeting and condensate behavior remain poorly understood. Using NUP98::KDM5A as a model, we integrate cellular imaging, *in vitro* reconstitution, and genomic analyses to dissect how chromatin engagement shapes condensate formation. We find that NUP98::KDM5A forms sub-diffraction-limited, gel-like condensates whose assembly is potentiated by binding to H3K4me3. This interaction creates a quantitative targeting mechanism in which, at the native expression level, condensates preferentially form at genomic loci with high local H3K4me3 density. Such local density-dependent recruitment explains selective enrichment at the leukemogenic HOX gene clusters, despite widespread presence of H3K4me3 across the genome. Analysis of single-cell sequencing data from patients further supports a correlation between local H3K4me3 density and transcriptional activation in NUP98::KDM5A-driven leukemia. Together, our findings reveal how activating chromatin marks and condensate-forming proteins synergize to generate specificity within euchromatin, offering a generalizable framework for understanding how chromatin-associated condensates interpret epigenetic landscapes.

## Introduction

Chromosomal translocations that generate NUP98-containing fusion proteins are a hallmark of multiple hematological disorders. These oncofusions are most prevalent in acute myeloid leukemia (AML) and are associated with poor prognosis (Bertrums et al., 2023; Gough et al., 2011; Tian et al., 2024; Umeda et al., 2025). While all NUP98 oncofusions share an N-terminal intrinsically disordered region (IDR) derived from NUP98, the C-terminal portion is derived from over 30 functionally diverse partner proteins (Michmerhuizen et al., 2020). The C-terminal fusion partners include those harboring a DNA- or chromatin-binding domain, allowing the oncofusions to localize to chromatin and alter gene expression. Several mechanisms of chromatin association have been described, including sequence-specific DNA-binding, such as NUP98::HOXA9 fusion, and specific histone mark recognition, exemplified by NUP98::KDM5A, NUP98-PHF23, and NUP98::KMT2A, all of which can bind trimethylated Lys4 in histone H3 (H3K4me3) through a PHD domain (Ahn et al., 2025; Calvo et al., 2002; Chandra et al., 2022; Gough et al., 2014; Wang et al., 2007, 2009).

NUP98 fusion oncoproteins have been shown to form condensates (Ahn et al., 2025; Chandra et al., 2022; Jevtic et al., 2022; Oka et al., 2023; Terlecki-Zaniewicz et al., 2021; Tripathi et al., 2023). These condensates may enrich general transcriptional machinery and transcriptional coactivators, including histone acetyltransferases, bromodomains, long noncoding RNAs, and the components of the menin-MLL complex, which promote sustained expression of leukemogenic genes (Ahn et al., 2025; Hamamoto et al., 2025; Heikamp et al., 2022, 2024; Michmerhuizen et al., 2025; Terlecki-Zaniewicz et al., 2021). Critically, the NUP98-derived region, which contains an FG-repeat sequence that mediates phase separation and condensate formation, is essential for leukemogenic transformation, as truncations and mutations of the phenylalanines in the FG repeats abrogate condensate formation and arrest the proliferation of hematopoietic stem and progenitor cells (Ahn et al., 2021, 2025; Chandra et al., 2022; Jevtic et al., 2022; Terlecki-Zaniewicz et al., 2021). In addition, chromatin binding is central to the leukemogenic potential of NUP98 fusion proteins, as mutational disruption of DNA recognition in NUP98::HOXA9 fusion and H3K4me3 binding by the PHD3 chromatin-binding domain in NUP98::KDM5A abolishes leukemogenic transformation (Ahn et al., 2021; Chandra et al., 2022; Wang et al., 2009). These findings indicate that both condensate formation and chromatin association of NUP98 oncofusions are required for leukemogenic transformation.

In normal cellular functions, several chromatin-associated condensates have been described, and they are associated with distinct chromatin states. For instance, the association between heterochromatin marks, like H3K9me3 and H3K27me3, and repressive factors, including HP1 and PRC1, leads to the formation of heterochromatin condensates, which compact chromatin and enrich silencing machinery to enforce a repressive chromatin state (Ackermann & Debelouchina, 2019; Deshpande et al., 2024; Larson et al., 2017; Plys et al., 2019; Sanulli et al., 2019; Strom et al., 2017; Tatavosian et al., 2019; Wang et al., 2019; Zhen et al., 2014). On the other hand, condensates associated with active, decompacted euchromatin are less well understood, with the exception of transcriptional condensates at super-enhancers that involve transcriptional coactivators and condensates at active gene arrays such as histone locus bodies (Sabari et al., 2018; Strom et al., 2024; Zhang et al., 2021). Oncofusions associated with the active-promoter histone mark H3K4me3, such as NUP98::KDM5A, offer disease model systems to probe the function of euchromatin-associated condensates.

Because HOXA9 is a leukemogenic transcription factor in itself, oncofusions with its DNA-binding domain enable targeting the NUP98::HOXA9 fusion to critical proleukemogenic genes. By contrast, H3K4me3 association could bring NUP98 fusions with chromatin readers to all active promoters across the genome. How these oncofusions bias the activation of leukemogenic genes remains an open question. Here, we address this key question using NUP98::KDM5A as our model system. Using a combination of cellular microscopy and *in vitro* reconstitution, we showed that NUP98::KDM5A forms gel-like condensates, and condensate formation is enhanced by H3K4me3 chromatin association. This association leads to a concentration-dependent selectivity of condensate targeting, with condensates preferentially forming at genomic loci with high local H3K4me3 mark density at pathologically relevant expression levels, such as the leukemogenic HOX gene clusters. We confirmed the correlation between gene activation and local H3K4me3 density by analyzing patient sequencing data. Our results not only provide mechanistic insights into the leukemogenesis process of NUP98 oncofusions involving chromatin readers, but also illustrate how functional specificity can be attained through the quantitative interplay between epigenetic landscapes and condensate-forming chromatin factors.

## Results

### NUP98::KDM5A forms condensates at H3K4me3-marked chromatin in a concentration-dependent manner

We sought to establish an expression system and to dissect the molecular determinants governing the cellular behavior of the oncogenic NUP98::KDM5A fusion protein. With this aim, we generated a series of Mammalian Toolkit-based plasmids (Fonseca et al., 2019) expressing fluorescently tagged NUP98::KDM5A and its variants and introduced them into cells via transient transfection, yielding cell populations with a wide range of expression levels in a single experiment (Chong et al., 2018, 2022). Live-cell imaging of U2OS cells revealed that expression of mEGFP-tagged NUP98::KDM5A leads to the formation of dense nuclear puncta, in contrast to mEGFP-tagged full-length KDM5A at similar expression levels (Figure 1 – Figure Supplement 1). However, the limited resolution of spinning-disk confocal microscopy hindered accurate segmentation of the puncta for quantitative analysis. Therefore, in our subsequent cell imaging experiments, we utilized the NSPARC scanning confocal microscope or the SoRa spinning disk confocal microscope, which have enhanced spatial resolutions (∼ 1.4x) to resolve neighboring NUP98::KDM5A puncta. For NUP98::KDM5A with mEGFP tagged on the C-terminus (NUP98::KDM5A-mEGFP), nuclear puncta almost all have sub-resolution-limit sizes (< 150 nm) across the expression level range (Figure 1A). Quantitative analysis of NSPARC images, followed by calibration of EGFP concentration to the microscope-acquired fluorescence signal (Figure 1 – Figure Supplement 2C and D), shows a threshold concentration of ∼50 nM for puncta detection (Figure 1C). As the overall nuclear concentration of NUP98::KDM5A (C_NUP98::KDM5A_) increases, the density of puncta rises, plateauing at ∼0.12 μm^-2^ in a given z-slice. When the concentration is above ∼1 μM, approximately 40% of the total fusion protein resides in these small puncta (Figure 1C). We estimated patient cellular concentration of NUP98::KDM5A to be ∼180 nM, which falls within the transition range (Figure 1 – Figure Supplement 3A,B; see Estimation of NUP98::KDM5A expression in patient cells in Materials and Methods). The small, dense nuclear puncta formed by NUP98::KDM5A are shaped by binding to H3K4me3 sites, which is supported by the observation that the chromatin-binding-deficient mutant (NUP98::KDM5Amut-mEGFP) forms large, spherical droplets that often coincide with nucleoli (Figure 1 – Figure Supplement 2A and B).

**Figure 1.**
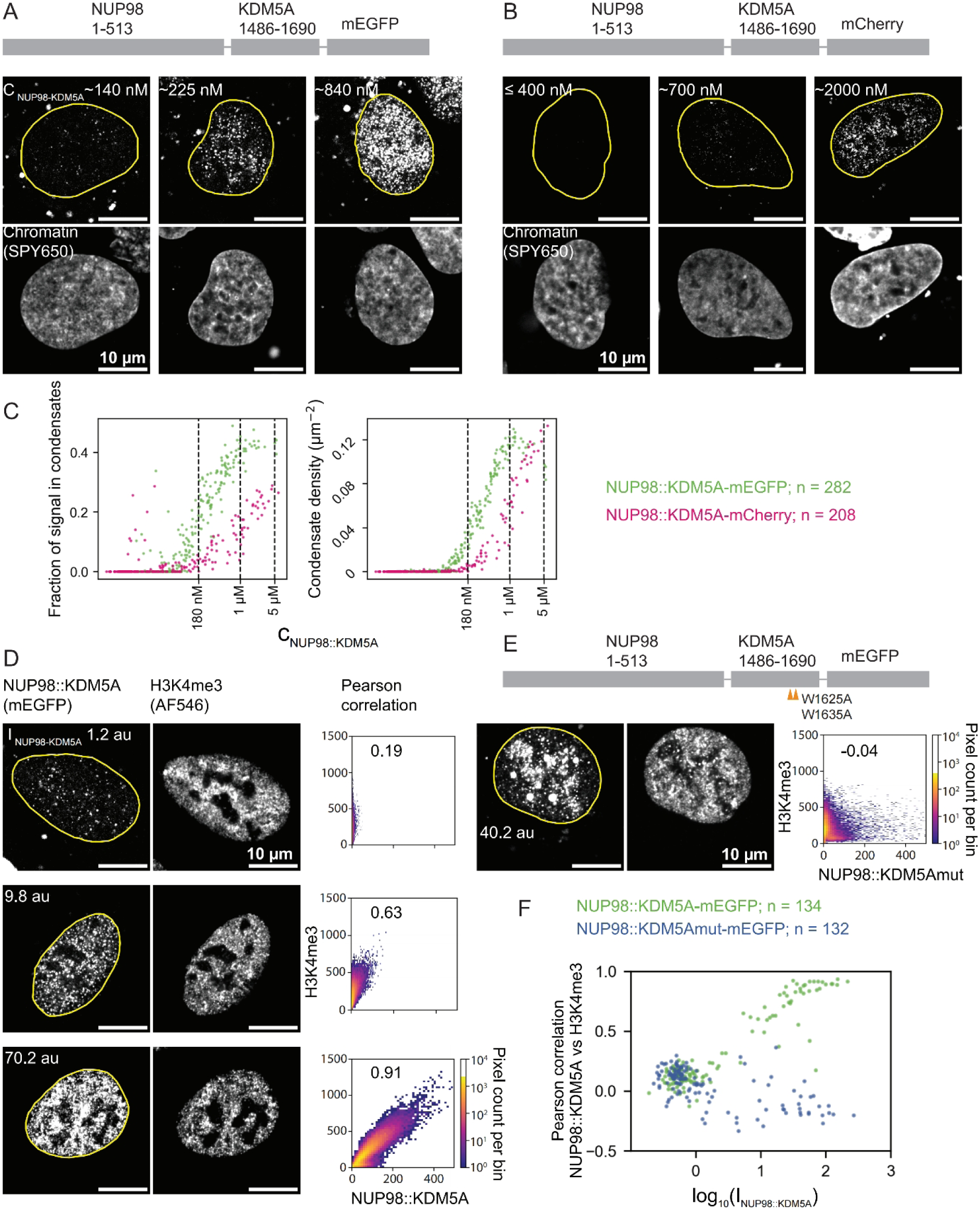
Concentration-dependent formation of H3K4me3-associated condensates by NUP98::KDM5A. **A.** Schematic of NUP98::KDM5A construct, with the N-terminal region of NUP98 (amino acids 1-513) fused to the C-terminal region of KDM5A (1486-1690), which includes the H3K4me3-binding PHD3 domain, and NSPARC images of NUP98::KDM5A-mEGFP in U2OS nuclei at different concentrations showing sub-resolution-limit puncta. **B.** Schematic of the NUP98::KDM5A-mCherry construct and NSPACR images in U2OS nuclei at different concentrations. **C.** Fraction of signal in condensates (fluorescent protein signal within condensates relative to total nuclear signal) and condensate density (number of condensates per µm^2^) as a function of nuclear protein concentration in individual cells (n = 282 cells for mEGFP tagging and n = 208 for mCherry tagging). **D.** Fixed U2OS nuclei expressing NUP98::KDM5A-mEGFP and immunostained for H3K4me3 with Alexa Fluor 546 (AF546), showing images with different mean nuclear mEGFP intensities (I_NUP98::KDM5A_), 2D histograms of pixel intensities, and the Pearson correlation, R, between the images of the two channels. **E.** Schematic of the chromatin-binding-deficient NUP98::KDM5A (W1625A, W1635A) construct, and the same set of analyses as in D. **F.** Image Pearson correlation, R, as a function of expression level, I_NUP98::KDM5A_, for U2OS cells expressing NUP98::KDM5A-mEGFP (n = 134 cells) or NUP98::KDM5Amut-mEGFP (n = 132).

To further test whether puncta formation involves phase separation in addition to chromatin binding, we replaced mEGFP with mCherry. While similar in size to mEGFP, mCherry has higher solubility and has previously been shown to promote the dissolution of biomolecular condensates (Brumbaugh-Reed et al., 2024; Mestrom et al., 2019). Indeed, compared to NUP98::KDM5A-mEGFP, NUP98::KDM5A-mCherry shows nuclear puncta formation at a higher concentration (∼200 nM), and a lower fraction of fusion protein molecules was localized to these puncta. NUP98::KDM5A-mCherry nuclear puncta have a similarly sub-resolution-limited size and plateau at a similar density (Figure 1B, C). The observation that NUP98::KDM5A puncta are formed by phase separation is further supported by the fact that the KDM5A-derived fragment of the oncofusion alone, which mediates chromatin-binding, does not form nuclear puncta when overexpressed in U2OS cells (Figure 1 – Figure Supplement 1A). Thus, we conclude that nuclear NUP98::KDM5A puncta are condensates formed through a combination of chromatin binding and phase separation.

The plateauing of NUP98::KDM5A condensate density at high concentration suggests saturation of H3K4me3 binding sites. To further characterize the occupancy of H3K4me3 sites by NUP9-KDM5A, we performed immunofluorescence of H3K4me3 and analyzed its image correlation with NUP98::KDM5A-mEGFP (Figure 1D). Pixel intensity histograms clearly show that, at low expression levels, NUP98::KDM5A-mEGFP condensates occupy a subset of H3K4me3 sites, whereas in cells with high expression, H3K4me3 sites fully colocalize with NUP98::KDM5A-mEGFP condensates. In contrast, the chromatin-binding-deficient NUP98::KDM5Amut-mEGFP shows complete anti-colocalization with H3K4me3 (Figure 1E). These trends are confirmed by the image Pearson correlation values as a function of NUP98 fusion expression inferred from mean nuclear mEGFP fluorescence intensity (I_NUP98::KDM5A_). The correlation between NUP98::KDM5A-mEGFP and H3K4me3 rises from zero to almost unity when NUP98::KDM5A-mEGFP expression exceeds a threshold, whereas that between NUP98::KDM5Amut-mEGFP and H3K4me3 remains low across the expression level range and has a negative trend (Figure 1F).

Taken together, our results indicate that NUP98::KDM5A forms nuclear, nanometer-sized condensates seeded at H3K4me3-marked sites through phase separation, and that both the fusion oncoprotein and its chromatin association determine the saturation concentration and biophysical properties of these condensates. As NUP98::KDM5A concentration increases, progressively more H3K4me3 sites become occupied until no additional sites remain available to support condensate formation.

### H3K4me3 binding enhances the formation of NUP98::KDM5A condensates in vitro

To quantitatively evaluate the interplay between NUP98::KDM5A and H3K4me3 chromatin in condensate formation, we used a reconstitution system that contains only purified components. To establish a concentration threshold for the formation of condensates *in vitro*, purified NUP98::KDM5A was labeled with the fluorescent dye Alexa Fluor 488 (AF488), and concentration-dependent condensate formation was monitored by confocal microscopy (Figure 2 – Figure Supplement 1A). Similar to other NUP98 fusions (Chandra et al., 2022), we found that at concentrations between 5.8 and 11.7 nM, full-length NUP98::KDM5A forms condensates in Phosphate Buffered Saline (PBS) with irregular, submicron-sized morphological features (Figure 2 – Figure Supplement 1A).

**Figure 2.**
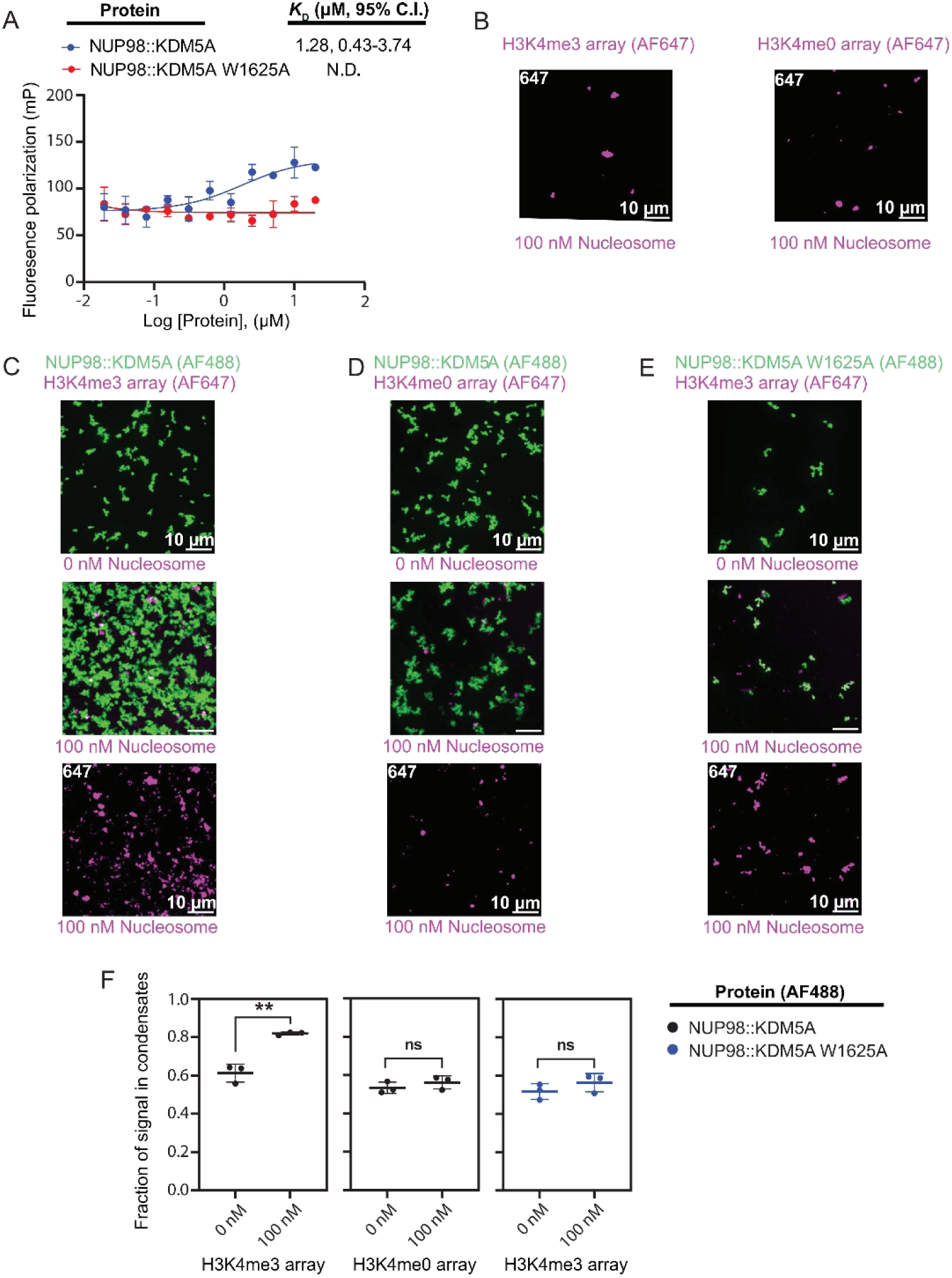
Binding of the PHD3 domain of NUP98–KDM5A to H3K4me3 promotes the formation of NUP98–KDM5A condensates. **A.** Fluorescence polarization (FP) assays to determine the *K*_D_ of NUP98::KDM5A WT or W1625A mutant for a 10-mer FAM-labeled H3K4me3 peptide (10 nM). Error bars represent standard deviations from technical triplicates. *K*_D_ values show means and 95% C.I.. N.D.: Not determined because no binding was observed. **B-E.** Representative confocal images of **B** AF647 labeled H3K4me3 or H3K4me0 arrays (100 nM nucleosomes), **C** AF488 labeled NUP98::KDM5A (800 nM) with AF647 H3K4me3 arrays (0 or 100 nM nucleosomes), **D** AF488 NUP98::KDM5A (800 nM) with AF647 H3K4me0 arrays (0 or 100 nM nucleosomes), or **E** AF488 NUP98::KDM5A W1625A (800 nM) with AF647 H3K4me3 arrays (0 or 100 nM nucleosomes). All images are maximum intensity projections (scale bar = 10 µm). **F:** The fraction of protein fluorescence signal in condensates relative to total signal in the image for the conditions in **C**-**E**. n = 3 technical replicates; ns: P > 0.05; *: P ≤ 0.05; **: P ≤ 0.01, parametric unpaired t-test.

The N-terminal FG-rich IDR of NUP98 has been previously shown to be essential for condensate formation (Ahn et al., 2021, 2025; Chandra et al., 2022; Jevtic et al., 2022; Terlecki-Zaniewicz et al., 2021). To evaluate the contributions of the sequence elements of the KDM5A fusion partner on the apparent saturation concentration (C_sat_), we generated three additional constructs: (1) NUP98::KDM5A(1601-1662), which retains only the PHD3 domain of the KDM5A fusion partner; (2) NUP98::KDM5A(1486-1600), where the KDM5A fusion partner is truncated at its C-terminus to remove the PHD3 domain, and (3) the NUP98 segment alone (Figure 2 – Figure Supplement 1A). While the PHD3-domain containing construct NUP98::KDM5A (1601-1662) exhibited a C_sat_ range similar to the full-length NUP98–KDM5A fusion (C_sat_: 5.8-11.7 nM), both the truncated fusion construct lacking the PHD3 domain and the NUP98 alone formed condensates at higher protein concentrations (C_sat_: 23.4-46.9 nM) (Figure 2 – Figure Supplement 1A). These findings suggest that, in addition to the previously characterized interactions between the FG domains of NUP98, the PHD3 domain of KDM5A significantly impacts the condensate behavior of NUP98::KDM5A.

We next evaluated the ability of the fusion protein to recognize the H3K4me3 modification by assessing binding to a fluorescently labeled H3K4me3 10mer peptide using a fluorescence polarization (FP)-based assay (Figure 2A). We also introduced an H3K4me3-binding-deficient PHD3 mutant into the fusion protein to modulate its ability to recognize the modification (Figure 2A). We found that the WT NUP98::KDM5A construct bound the H3K4me3 peptide with a dissociation constant (*K*_D_) of 1.28 µM (95% CI, 0.43-3.74 µM), which is consistent with *K*_D_ values reported for the isolated PHD3 domain of KDM5A (Figure 2A) (Zhang et al., 2022). In contrast, the chromatin-binding-deficient mutant NUP98::KDM5A W1625A exhibited no measurable binding with increasing protein concentrations of up to 20 µM (Figure 2A). These results indicate that the PHD3 domain in the NUP98::KDM5A fusion protein retains the ability to recognize the H3K4me3 mark in the context of the histone H3 peptide.

We then sought to examine the effect of NUP98::KDM5A-nucleosome interactions on the behavior of NUP98::KDM5A condensates. In these experiments, we carried out *in vitro* phase separation assays using either chicken erythrocyte polynucleosomes as a substrate with a heterogeneous pattern of histone modifications, or reconstituted nucleosomal arrays with a uniform pattern of histone modifications (H3K4me3 or -me0). In the presence of chicken erythrocyte polynucleosomes, there is a roughly 15% increase in the fraction of NUP98::KDM5A incorporated into condensates relative to reactions lacking chicken erythrocyte polynucleosomes (Figure 2 - Figure Supplement 1B and D). In contrast, the addition of chicken erythrocyte polynucleosomes to NUP98::KDM5A W1625A had no significant effect on the fraction of molecules partitioned into condensates (Figure 2 - Figure Supplement 1C and D). These observations imply chromatin binding enhances NUP98::KDM5A condensate formation.

It is well characterized that FG-repeat IDRs from NUP98 and other nucleoporins can form gel-like condensates, which are important for controlling nuclear transport by the nuclear pore (Schmidt & Görlich, 2015). The irregular morphology of reconstituted NUP98::KDM5A condensates also indicates their gel-like states. Indeed, fluorescence recovery after photobleaching (FRAP) measurements of NUP98::KDM5A condensates show little recovery both in the absence of polynucleosomes (mobile fraction, M_F_ = 0.043) and with the addition of chicken erythrocyte polynucleosomes (M_F_ = 0.072) (Figure 2 - Figure Supplement 2A). Consistent with reconstituted condensates, live cell FRAP measurements of NUP98::KDM5A-mEGFP also show a low mobile fraction (M_F_ = 0.085) (Figure 2 - Figure Supplement 2A). Interestingly, switching the mEGFP tag to N-terminus dramatically increases the fluidity of NUP98::KDM5A condensates in living cells (M_F_ = 0.48 and half-time of recovery T_1/2_ = 1.9 sec) (Figure 2 – Figure Supplement 2A), despite a similarly small, punctate appearance of the condensates, comparable fraction of signal in condensates and condensate density regardless of the position of the mEGFP tag (Figure 2 – Figure Supplement 2B-D). A possible explanation for this effect is the insolubility of the mEGFP tag in fluorescence protein-repeat condensates (Frey et al., 2018), which could disrupt gel networks when these two components are immediately adjacent in the N-terminal tagged construct. This observation may explain the variability of fluidity reported for NUP98 oncofusion condensates (Chandra et al., 2022; Ibáñez De Opakua et al., 2022; Schmidt & Görlich, 2015). It also provides another vivid example of how tagging could affect the biophysical properties of biomolecular condensates (Fatti et al., 2025; Zhou et al., 2025). Based on these findings, we used C-terminal mEGFP tagging for all our live-cell experiments.

To further interrogate the role of chromatin ligands in the formation and the morphology of NUP98::KDM5A condensates, we evaluated the influence of substrates with a homogeneous pattern of histone modifications. For this purpose, we prepared arrays with histone octamers containing either unmodified (H3K4me0) or trimethylated (H3K4me3) H3 reconstituted onto AF647-labeled DNA consisting of a previously characterized 3.5 kb DNA sequence from the 5’ end of the mouse gene *Cyp3a11*, which contains approximately 16 nucleosomes (Luger et al., 1999; Moore et al., 2025). Consistent with previous studies, we found that nucleosomal arrays, in the absence of NUP98::KDM5A, crowding agents, or added divalent cations, can form condensates under physiological concentrations of monovalent salts (Moore et al. 2025, Gibson et al. 2019) (Figure 2B) (Figure 2 – Figure Supplement 3A-C). The reconstituted H3K4me3 or H3K4me0 nucleosome arrays were then mixed with AF488-labeled NUP98::KDM5A at a physiological salt concentration (136.9 mM NaCl, 2.68 mM KCl), and condensates were monitored by confocal microscopy (Figure 2C, D, Figure 2 – Figure Supplement 3A, B). Our results show that *in vitro* mixtures of NUP98::KDM5A with H3K4me3 or H3K4me0 nucleosomes showed a similar degree of colocalization, with the highest degree of colocalization at 12.5 nM nucleosomes (Figure 2C, D) (Figure 2 – Figure Supplement 3A, B). We also observed that mixtures of NUP98::KDM5A W1625A with H3K4me3 nucleosomes (Figure 2E) (Figure 2 – Figure Supplement 3C) exhibited a similar degree of colocalization as mixtures of NUP98::KDM5A with H3K4me3 (Figure 2C) (Figure 2 – Figure Supplement 3A) or H3K4me0 nucleosomes (Figure 2D) (Figure 2 – Figure Supplement 3B). Interestingly, we found that the presence of H3K4me3 nucleosomes led to approximately a 20% increase in the fraction of NUP98–KDM5A signal in condensates, which appear irregularly shaped, highly interconnected, and gel-like (Figure 2C, F). In contrast, no significant changes were observed in the fraction of signal incorporated into condensates when the WT NUP98::KDM5A was incubated with the H3K4me0 nucleosome array (Figure 2D, F), or when the NUP98–KDM5A W1625A mutant protein was incubated with the H3K4me3 nucleosome array (Figure 2E, F). Together, our findings reveal that the interaction between the PHD3 domain in the NUP98::KDM5A fusion protein and the H3K4me3 mark is a critical modulator of the spatial organization and behavior of NUP98::KDM5A condensates.

### NUP98::KDM5A preferentially forms condensates at H3K4me3-enriched loci

A survey of publicly available H3K4me3 ChIP-seq data revealed that HOX clusters, which are key regulators of developmental processes and contain prominent leukemogenic genes such as HOXA9, are among the most densely H3K4-trimethylated regions across many cell types. We thus tested whether high local H3K4me3 density leads to preferential formation of NUP98::KDM5A condensates. We performed fluorescence *in situ* hybridization (FISH) combined with immunofluorescence in HEK293 cells to quantify the colocalization between condensates formed by C-terminally FLAG-tagged NUP98::KDM5A (NUP98::KDM5A-FLAG) and various genomic loci, including the HOXA/B/D clusters with high H3K4me3 density, inactive gene TNF but with H3K4me3 from nearby genes, as well as the inactive gene MUC4 and the active gene MYC, both having very low total H3K4me3 within a ±50 kb window (Figure 3A). For each locus, we used two differently colored sets of FISH probes targeting contiguous sequences to identify specific signals as paired spots in NSPARC enhanced-resolution 3D images (centroid distance < 300 nm). We then quantified NUP98::KDM5A-FLAG condensate association by measuring the local NUP98::KDM5A signal, defined as the maximum local intensity (within 200 nm 3D radius from the loci centroid) minus the neighboring background (the 20th-percentile intensity within 800 nm 3D radius) (Figure 3B). Figure 3C shows this local NUP98::KDM5A signal for each identified locus in the FISH image as a function of the NUP98::KDM5A expression level, which is represented by the mean nuclear fluorescence signal (I_NUP98::KDM5A_).

**Figure 3.**
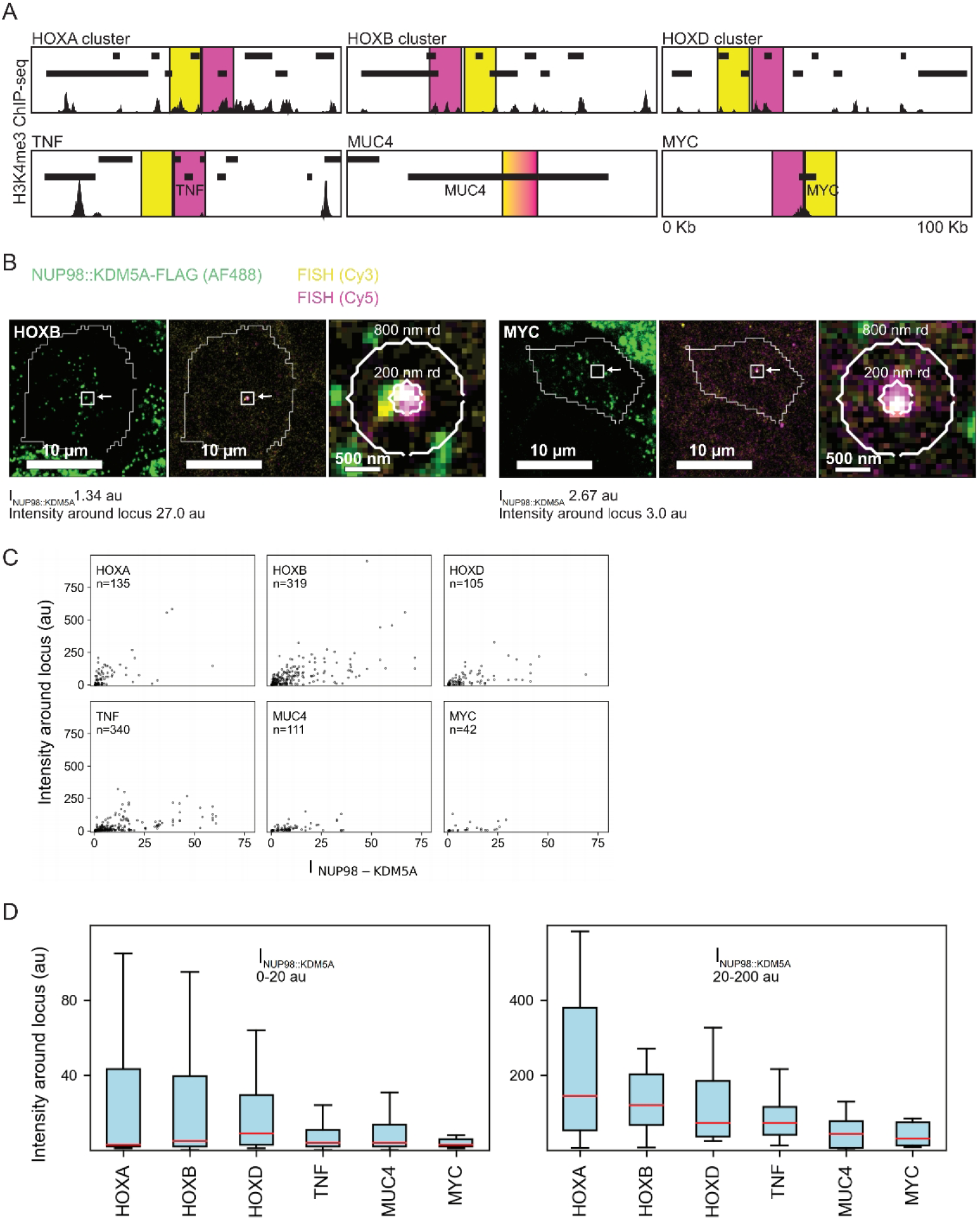
NUP98::KDM5A condensate formation at loci with varying H3K4me3 density. **A.** Schematic of the 100 kb genomic region surrounding each FISH-labeled locus. Bars indicate genes. Black peaks at the bottom show H3K4me3 ChIP-seq signal in HEK293 cells. FISH probe targeted regions are marked in magenta (Cy5-labeled probes) and yellow (Cy3-labeled probes). **B.** Images of combined FISH (magenta and yellow) and NUP98::KDM5A immunofluorescence (green) in HEK293 cells for the HOXB and MYC loci, showing a single z-slice with white lines marking the nucleus based on DAPI signal segmentation. Circles in the zoomed-in views of the boxed regions mark the radii for maximum and background signal calculation. **C.** Local NUP98::KDM5A signal (intensity around the locus) across expression levels as measured by mean NUP98::KDM5A nuclear intensity, I_NUP98::KDM5A_, (HOXA, n = 135 loci; HOXB, n = 319; HOXD, n = 105; TNF, n = 340; MUC4, n = 111; MYC, n = 42). **D.** Boxplots of local NUP98::KDM5A signal for cells with low (I_NUP98::KDM5A_ = 0 ∼ 20 au) and high (I_NUP98::KDM5A_ = 20 ∼ 200 au) expression levels. The median is shown in red. The box extends from the first to the third quartile, and whiskers extend to the most extreme data points located within 1.5x the interquartile range.

All six scatter plots in Figure 3C show a general trend that a higher expression level leads to more loci exhibiting a high local NUP98::KDM5A signal, i.e., association with a NUP98::KDM5A condensate, which is consistent with the trend shown in Figure 1D. At a low expression level, on the other hand, a substantial fraction of HOXA/B/D loci still have high local NUP98::KDM5A signal, whereas the vast majority of TNF, MUC4, and MYC loci show low local NUP98::KDM5A signal. This difference is more clearly shown in the box plots (Figure 3D) by dividing cells into low-expression (I_NUP98::KDM5A_ = 0 ∼ 20 au) and high-expression (I_NUP98::KDM5A_ = 20 ∼ 200 au) groups. As a control, we performed the same analysis on the DAPI DNA stain channel in the same images (Figure 3 - Figure Supplement 1), which show low values for all local DAPI signals and flat trends. Taken together, these results indicate that, although a high expression level of NUP98::KDM5A can drive condensate formation at all gene loci, a low expression level leads to preferential formation of condensate at loci with high H3K4me3 density, such as the leukemogenic HOX gene clusters.

### The local density of H3K4me3 correlates with expression upregulation in patient cells

Having established that NUP98::KDM5A forms condensates at H3K4me3-rich loci in a concentration-dependent manner in HEK293 cells, we next examined how NUP98::KDM5A affects gene expression at potential target loci under patient-relevant NUP98::KDM5A expression levels and conditions. We analyzed published single-cell RNA-seq and H3K4me3 CUT&RUN datasets from primary leukemia patients harboring the NUP98::KDM5A oncofusion (Umeda et al., 2025). We analyzed each patient separately, i.e., one with acute megakaryoblastic leukemia (Patient 1; AMKL) and one patient with acute lymphoblastic leukemia (Patient 2; ALL), as the transcriptome profiles of these leukemias are distinct (Umeda et al., 2025).

First, we assessed fold changes in gene expression in patients compared to age- and cell-matched healthy individuals. A separate comparison was made for each patient due to the inherent heterogeneity of leukemia transcriptomes. Loci were defined as 100 kb windows (±50 kb) centered on transcription start sites (TSSs) (Figure 4 - Figure Supplement 1A), as previous studies have described broad 60-100 kb as critical descriptors of cell identity (Benayoun et al., 2014). Volcano plots indicate that, in both patients, more genes are upregulated than downregulated compared with healthy individuals. The upregulation is consistent with the proposed gene-activating function of NUP98::KDM5A condensates, whereas the downregulation likely reflects complex interactions within gene regulatory networks that lead to repression at specific loci (Figure 4A). Genes at high H3K4me3 loci exhibit elevated mean expression and are expressed in a greater proportion of cells in patients than in healthy donors, despite inter-donor variability (Figure 4B).

**Figure 4.**
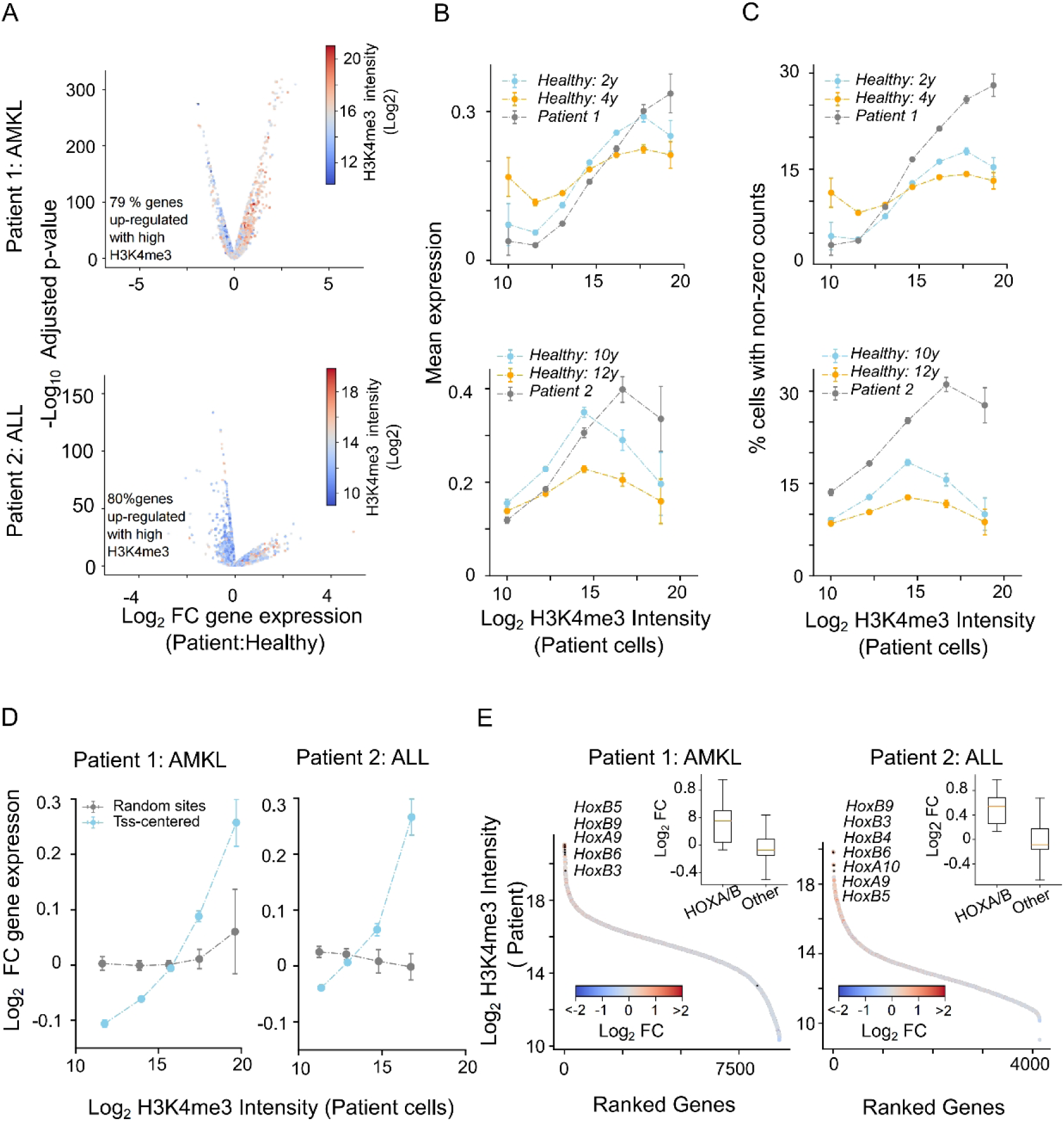
Degree of expression correlates with H3K4me3 intensity for dysregulated genes in NUP98::KDM5A leukemia patients. **A.** Volcano plots show changes in gene expression in patients compared to healthy donors, with color encoding H3K4me3 intensity. Rows correspond to patients. **B-C.** Donor-specific trends in H3K4me3-associated gene expression in patients compared with their matched healthy controls. **B.** Mean gene expression across genes within each H3K4me3 intensity bin at a 100-kbp window. The mean represents the average gene-specific mean expression across cells within each bin. Similarly, **C.** proportion of cells with detectable gene expression within the same bins. Error bars: SEM. **D.** Correlation plot of change in gene-expression with H3K4me3 intensity in patients with genes binned by methylation signal and expression fold change. Blue: intensities computed across TSS-centered windows, and Gray: across random sites for comparable window size. Error bars: SEM. **E.** Rank-ordered H3K4me3 intensity plot with color representing the fold-change in gene expression. Top-ranked HOXA and HOXB protein-coding genes with log2 fold change ≥ 0.5 are indicated. Inset: boxplot represents the fold-change in gene-expression for HOXA/B coding genes compared to all the other genes in the ranked set (Patient 1: n = 16 for HOXA/B and 9205 for the rest; Patient 2: n = 12 for HOXA/B and 4125 for the rest).

Next, we examined how H3K4me3 levels at loci correlated with fold changes in gene expression in the two patients in our study. In both patients, the majority of dysregulated genes were heavily H3K4me3-marked (Figure 4A). We observed a clear positive correlation, with the most highly H3K4me3-marked loci also being the most upregulated (Figure 4B). To better visualize the relationship between the presence of H3K4me3 and gene expression changes at target loci, we ranked TSS-centered loci according to their H3K4me3 content (for all genes filtered by FDR-adjusted p-value ≤ 0.05; rank 1 representing the highest intensity) and quantified their fold-change in expression relative to healthy individuals. This analysis revealed a clear positive correlation, with the most highly H3K4me3-marked loci (left) exhibiting the strongest upregulation. Notably, the loci with the highest H3K4me3 levels corresponded to the HOXA and HOXB clusters. Furthermore, fold changes at the HOXA and HOXB clusters exceeded those of other upregulated genes (Figure 4C). Notably, plots from the two patients reveal distinct transcriptional programs in NUP98::KDM5A-driven cancers, likely reflecting differences in developmental stage and unique passenger mutations in the precursor cells.

Taken together, our results show that the most dysregulated genes in the patients are also heavily H3K4me3-marked, predominantly up-regulated, and associated with leukemogenesis. Among these dysregulated genes, the *HOXA* and *HOXB* clusters stand out, exhibiting the highest H3K4me3 levels in patient cells, consistent with their well-established role in NUP98::KDM5A-driven leukemia (Noort et al., 2021). Importantly, to account for the activating effect of H3K4me3 itself, our analysis explicitly uses fold-change between patient cells and healthy controls as the readout. This correlative trend, together with the observation that gene expression saturates over a length scale of approximately ±50 kb (Figure 4 – Figure Supplement 1), further supports selective enhancement of condensate formation in the presence of the NUP98::KDM5A oncofusion until available H3K4me3 sites become saturated.

## Discussion

Our results revealed key interactions associated with the formation of NUP98::KDM5A condensates in euchromatin loci relevant for leukemogenic transformation. Specifically, we showed that the formation of NUP98::KDM5A condensates involves a combination of chromatin binding and phase separation, and that the interplay between these distinct types of interactions enhances NUP98::KDM5A condensation. Additionally, our findings address long-standing questions about how condensates formed by NUP98 fusions with chromatin readers are targeted to specific chromatin loci. We envision that a similar mechanism targets other NUP98 fusions containing H3K4me3-recognizing reader domains (e.g., NUP98::PHF23) to leukemogenic loci, complementing direct targeting of the *HOX* genes through DNA sequence recognition in fusions containing DNA-binding domains (e.g., NUP98::HOXA9). Despite being a pathological protein, condensates formed by NUP98::KDM5A show several features that could be generalizable to physiological chromatin-associated condensates, e.g., those formed by transcription factors.

First, NUP98::KDM5A condensates show clear gel-like properties with low exchange dynamics, contrasting earlier impressions in the biomolecular condensate field that liquid-like fluidity or fast exchange dynamics are required for their functions. In fact, it is now recognized that certain chromatin-protein co-condensates can be strongly solid-like, such as those formed in olfactory neurons to enable the silencing of all but one olfactory receptor gene (Pulupa et al., 2025). Interestingly, rather than silencing, NUP98::KDM5A condensate formation actually activates targeted genes. The gel-like properties of NUP98::KDM5A condensates may enable this role not only by enriching activating factors, but also by excluding repressive factors, analogous to the selectivity of gel-like condensates in the nuclear pore formed by the same FG-repeat IDRs from NUP98 and other nucleoporins (Hulsmann et al., 2012). This exclusion could be easily overlooked because it is more difficult to measure by proteomics methods than enrichment.

Second, NUP98::KDM5A condensates are dispersed into small, sub-diffraction-limit sizes that are associated with the cognate H3K4me3 domains. Interestingly, in the cell nucleus, this small size does not rely on its gel-like properties, because the N-terminally tagged mEGFP-NUP98::KDM5A shows fast FRAP recovery but similar morphologies to the gel-like C-terminally tagged NUP98::KDM5A-mEGFP. Instead, chromatin binding is key to this appearance, because nuclear condensates formed by the chromatin-binding-deficient mutant are substantially larger (lower condensate density, but similar fraction of proteins in condensates). It could be imagined that binding to specific chromatin sites could also enable other transcription factors to form sub-diffraction-limit condensates (or clusters) that are challenging to detect by conventional fluorescence microscopy techniques.

Finally, the formation of small, chromatin-associated NUP98::KDM5A condensates at specific loci depends on both expression level and binding-site density. For instance, it remains poorly understood how chromatin readers recognize critical target loci. Although histone marks are known to mediate the association of epigenetic readers with chromatin, these marks are often widespread across the genome, and the sources of binding specificity remain unclear (Lukauskas et al., 2024; Yun et al., 2011). Binding of epigenetic readers to specific loci is frequently the first step in feedback loops that influence gene expression and ultimately cell-fate decisions (Atlasi & Stunnenberg, 2017; Blackledge & Klose, 2021; Meissner, 2010). Here, we show that chromatin association at specific loci emerges from the combined effects of NUP98::KDM5A expression level and local H3K4me3 density. At low expression levels, NUP98::KDM5A preferentially associates with highly H3K4me3-marked loci, whereas this preference diminishes as expression levels increase. These observations indicate a high degree of specificity of NUP98::KDM5A for loci enriched in the H3K4me3 mark. Furthermore, this suggests a potential general mechanism underlying transcription factor and epigenetic reader specificity and highlights the importance of tightly controlling the expression levels of chromatin-associated factors, as the abundance of these proteins serves as a critical regulatory knob for chromatin binding and the activation of downstream gene expression programs.

Study limitations

Cellular experiments in this study were performed using transient transfection of WT NUP98::KDM5A and its mutants in HEK293 and U2OS cells. While these non-hematopoietic cell models limit physiological relevance, they allowed for controlled expression levels in the absence of endogenous oncofusion and the ease of imaging, enabling investigation of the biophysical principles of condensate formation, the main objective of our study. Similarly, our study focuses on the biophysical mechanism of chromatin targeting by the NUP98::KDM5A oncofusion, with functional consequences inferred from patient-derived datasets rather than defined perturbation experiments.

## Materials and Methods

### DNA constructs for cell expression

We used the Mammalian Toolkit (MTK) system (Fonseca et al., 2019) to generate constructs expressing variants of the NUP98::KDM5A oncofusion in tissue culture. MTK enables Golden Gate–based assembly of modular parts into complete transcriptional units for expression via transient transfection. First, we generated a library of modular parts (NUP98 fragment, KDM5A-derived fragments, fluorescent proteins) either by PCR amplification or IDT DNA synthesis, ensuring that none of the sequences contained Esp3I or BsaI recognition sites. This “domestication” step is required because Esp3I and BsaI are the type IIS restriction enzymes used for cloning modular parts and for assembling transcriptional units, respectively. Esp3I digestion enables insertion of PCR-amplified sequences into domesticated parts. In contrast, BsaI digestion of the domesticated parts generates complementary sticky ends—designated 3a, 3b, and 4a—that direct ordered assembly of the transcriptional unit into the MTK acceptor expression vector (MTK0). The acceptor vector we used carries a pGK promoter, polyadenylation tail, and ampicillin resistance cassette. During the Golden Gate assembly, the 3a overhang anneals to its corresponding site in the acceptor vector, the 3b part ligates between the 3a and 4a overhangs, and the reaction closes with the 4a overhang, producing a correctly assembled circular expression vector suitable for expression of NUP98::KDM5A upon transient transfection. A list of acceptor vectors used in this study is provided in Supplementary Table 1. Construct sequences are provided in folders with the suffix _DNASequences in Dryad repository. Plasmids were purified using the BioBasic EZ-10 Spin Column Plasmid DNA Mini-Preps Kit (#C861B24). Cloning was performed in NEB Stable competent E.coli (NEB; C3040H).

The NUP98::KDM5A fusion sequence was derived from the cDNA sequences retrieved from NCBI Nucleotide (GenBank) for NUP98 isoform 1 (accession: NM_016320.5) and KDM5A (accession: NM_001042603.3). The fusion junction was selected as indicated in Figure 3 of (Van Zutven et al., 2006), incorporating amino acids 1-513 from NUP98 and 1486-1690 from KDM5A into the oncofusion. Synonymous nucleotide substitutions were introduced to the constructs to ensure compatibility with MTK Golden Gate cloning. Plasmid sequence files for plasmids used in mammalian cell expression experiments are stored in Figure1_DNASequences and Figure3_DNASequences folders in the Dryad repository.

### Cell lines and tissue culture

For experiments in the results section “NUP98::KDM5A forms condensates at H3K4me3-marked chromatin in a concentration-dependent manner”, we used U2OS cells (human osteosarcoma) purchased from ATCC (Catalog Number 50-238-4124). For results sections “NUP98::KDM5A preferentially forms condensates at H3K4me3-enriched loci” we used HEK 293 FT cells (human embryonic kidney) purchased from Invitrogen (Catalog Number R70007). Cell lines were maintained under standard culture conditions in standard T-25 or T-75 tissue culture-treated surface flasks in Dulbecco’s Modified Eagle Medium (DMEM; Gibco #10566024) supplemented with 10% Refiltered Fetal Bovine Serum (FBS; UCSF Media Production #CCFAP004) and 1% Penicillin-Streptomycin (11 mg/mL; UCSF Media Production #CCFGK004). Cultures were maintained in a mammalian cell culture humidified incubator at 37 °C adjusted to 5% CO_2_. Cells were re-plated every 3-5 days after one Phosphate Buffered Saline (PBS) (Gibco #20012027) wash and 5 min enzymatic cell detachment using Trypsine-EDTA 0.25% in Saline A (UCSF Media Production #CCFGP003), followed by gentle resuspension in culture media.

### *In vitro* reconstituted condensates: DNA constructs, protein expression, and purification

pET28a-based expression vectors were generated to produce the full-length NUP98::KDM5A oncofusion and other C-terminally truncated variants of NUP98::KDM5A (supplementary table 2). All expression constructs were generated by fast cloning using synthetic geneblock fragments (IDT) that were codon optimized for expression in BL21 (DE3) *E. coli* cells (C. Li et al., 2011). Plasmid sequence files for plasmids used in bacterial protein purification experiments are stored in Figure2_DNASequences folder in the Dryad repository.

### *In vitro* reconstituted condensates: Protein expression and purification

The protein expression and purification strategies for NUP98::KDM5A and other NUP98-containing constructs were carried out as previously described for the related NUP98-HOXA9 oncofusion and the NUP98 segment alone (Chandra et al., 2022; Ibáñez De Opakua et al., 2022; Schmidt & Görlich, 2015). All proteins used in this study were expressed in BL21(DE3) competent *E. coli* cells (NEB) that were transformed with pET28a-based plasmids and grown at 37 °C to an optical density at 600 nm (OD600) of approximately 0.8 in Luria-Bertani (LB) medium supplemented with Kanamycin. At an OD600 of ∼0.8, protein expression was induced by the addition of 1 mM IPTG (GoldBio), and the bacterial cultures were incubated at 37 °C for 4 hours. For protein purification, 8 liters of bacterial cultures were harvested by centrifugation (6,000 x g for 15 minutes at 4 °C). Bacterial cells were resuspended in Buffer A (20 mM Tris, 500 mM NaCl, 1 mM (tris(2-carboxyethyl)phosphine) (TCEP), 0.1 % Triton X-100, pH 7.4) and lysed by sonication on ice. The cell lysates were clarified by centrifugation at 30,000 x g at 4 °C and the supernatant was discarded as all the constructs of interest were found in the insoluble fraction. The inclusion bodies obtained were resuspended in extraction buffer (Buffer A supplemented with 6 M guanidine hydrochloride), followed by sonication and centrifugation. After the second centrifugation step, the supernatant of the inclusion bodies dissolved in extraction buffer was loaded onto a nickel-nitrilotriacetic acid (Ni-NTA) column (Thermo Fisher Scientific). The bound proteins were washed with 5 column volumes of extraction buffer containing 50 mM Imidazole and eluted with extraction buffer containing 500 mM Imidazole. Fractions containing the proteins of interest were concentrated and purified by size exclusion chromatography using a Superdex 200 16/600 column that was equilibrated with extraction buffer. High purity fractions were identified by SDS-PAGE, concentrated by ultracentrifugation using a 30 kDa molecular weight cutoff membrane, flash frozen and stored at -80 °C. Protein concentrations were determined using the following extinction coefficients at 280 nm: NUP98 (1-513)-KDM5A (1486-1690): 27430 M^-1^cm^-1^, NUP98 (1-513)-KDM5A (1601-1662): 18950 M^-1^cm^-1^, NUP98 (1-513)-KDM5A (1486-1600): 12950 M^-1^cm^-1^, and NUP98 (1-513): 5960 M^-1^cm^-1^.

### *In vitro* reconstituted condensates: AF488 labelling of proteins

All proteins were labeled using Alexa Fluor 488 TFP ester reactive dyes according to the manufacturer’s protocol. Briefly, proteins were diluted to 1 mg/mL using PBS (136.9 mM NaCl, 2.68 mM KCl, 10 mM Na_2_HPO_4_ 1.7 mM KH_2_PO_4_, pH 7.4) and then mixed 10:1 with 1 M sodium bicarbonate. The labelling reactions were carried out at a dye:protein molar ratio of 10:1. The reaction mixtures were then incubated for 15 minutes at room temperature. To remove any unbound dye from the protein solution, the reaction mixtures were dialyzed for 16 hours at 4 °C against the extraction buffer at 5,000 times the volume of the sample using a 2 kDa molecular weight cutoff membrane. For labeled proteins, the concentration and degree of labelling are based on extinction coefficients at 280 nm and were determined by measuring the absorbance at 280 nm and 494 nm.

### *In vitro* reconstituted condensates: Fluorescence polarization assay

The binding of unlabeled NUP98::KDM5A or NUP98::KDM5A W1625A to a H3K4me3 10mer peptide was measured either by direct or competition-based fluorescence polarization assays (FP). For the direct FP binding assays, increasing protein concentrations of up to 20 µM were incubated for 30 minutes at room temperature with 10 nM of a C-terminal fluorescently labeled H3K4me3 peptide. To determine the dissociation constant (K_d_) for the H3K4me3 peptide, the raw data from direct FP measurements were fitted to a one site-total nonlinear regression curve fit equation (GraphPad Prism 9.5.1, GraphPad Software, La Jolla, CA). For competition-based FP assays, 2 µM NUP98::KDM5A or MBP-PHD3 (KDM5A residues 1601-1662) was incubated for 30 minutes at room temperature with 10 nM of C-terminal fluorescently labeled H3K4me3 peptide in the presence of increasing concentrations of unlabeled competitor peptides. IC_50_ values were determined by fitting the raw data from competition-based FP measurements to an inhibitor versus response (three parameters) nonlinear regression curve fit equation (GraphPad Prism 9.5.1, GraphPad Software, La Jolla, CA). Direct or competition-based FP assays were carried out in 1X PBS (136.9 mM NaCl, 2.68 mM KCl, 10 mM Na_2_HPO_4_, 1.7 mM KH_2_PO_4_, pH 7.4) supplemented with 2.5 mM DTT and performed in triplicates.

### Nucleosome assembly and *in vitro* phase separation assays: Generation of AF647 labeled H3K4me3 or H3K4me0 nucleosomes

DNA was purified, labeled and assembled onto histone octamers as previously described (Luger et al., 1999; Moore et al., 2025). In brief, a 3.5 kb insert consisting of 3.2 kb from the 5′ end of the mouse *Cyp3a11* gene, 300 bp of plasmid DNA, and a 5′ overhang, was purified by size-exclusion chromatography and labeled with Alexa Fluor 647–aha-dCTP (Thermo Fisher Scientific) using the Klenow fragment (Moore et al. 2025). Unmodified (H3K4me0) or histone H3 lysine 4 trimethylated (H3K4me3) histone octamers (two each of human histones H2A, H2B, H3, and H4), made from recombinant histones expressed in *E. coli*, were purchased from EpiCypher. AF647 labeled DNA was assembled onto H3K4me0 or H3K4me3 histone octamers using a salt gradient dialysis at a histone octamer:DNA molar ratio of 17.7:1 (Luger et al. 1999). Following assembly, samples were dialyzed against TCS buffer (20 mM Tris-HCl, pH 7.5, 1 mM EDTA, 1 mM DTT) and stored at 4 °C. DNA concentration was determined after assembly via Nanodrop and the molecular weight of the double-stranded sequence, along with the histone octamer:DNA molar ratio, was used to calculate the nucleosome concentrations. Nucleosome assembly was assessed by protection of DNA from enzymatic digestion by Micrococcal nuclease.

### Nucleosome assembly and *in vitro* phase separation assays: Preparation of microscopy plates

384-well glass-bottom SenoPlates (Greiner Bio-One) were mPEGylated and passivated with Bovine serum albumin (BSA) as previously described (Hihara et al. 2012). Briefly, microscopy plates were washed and incubated with 2% Hellmanex for 1 hour at room temperature. After several washes with H_2_O, wells were incubated with 0.5 M NaOH for 30 minutes and then rinsed with H_2_O. For mPEG passivation, wells were incubated overnight at 4 °C with a solution of 5K mPEG-silane (Laysan Bio) at 20 mg/mL dissolved in 95% Ethanol. The wells were washed twice with 95% Ethanol, followed by three washes with H_2_O, prior to the addition of a 100 mg/mL solution of BSA dissolved in H_2_O. To remove the BSA solution, wells were washed six times with H_2_O. After three additional washes with phase separation assay buffer (1X PBS (136.9 mM NaCl, 2.68 mM KCl, 10 mM Na_2_HPO_4_, 1.7 mM KH_2_PO_4_, pH 7.4) supplemented with 2.5 mM DTT), the plates were ready for use.

### Nucleosome assembly and *in vitro* phase separation assays: *In vitro* phase separation assay

*In vitro* assays were prepared as previously described with minor modifications and protein concentrations expressed as mixtures of AF488 labeled:unlabeled proteins at a ratio 1:10 mixtures (Chandra et al. 2022, Ruan et al. 2025). Briefly, unlabeled stocks were first diluted to 300 µM in extraction buffer and then gradually diluted to 1.5 M guanidine hydrochloride with H2O before mixing with labeled samples to make a 37.5X mixture at 1:10 (labeled to unlabeled). The 1:10 protein mixture was further diluted to 4.75X (0.1M guanidine hydrochloride) in H2O, and serial dilutions in the same solvent were carried out. The 1:10 protein mixtures were then mixed with phase separation assay buffer (1X PBS (136.9 mM NaCl, 2.68 mM KCl, 10 mM Na_2_HPO_4_, 1.7 mM KCl, pH 7.4) supplemented with 2.5 mM DTT) to achieve the final concentrations and negligible levels of guanidine hydrochloride (0.02 M).

For all *in vitro* phase separation reactions, samples were incubated at room temperature overnight before imaging in mPEG/BSA passivated 384-well SensoPlates (Greiner Bio-One). Phase separation reactions with increasing concentrations of proteins (0-750 nM, Supplementary Figure 2A) were prepared as described above, where 2-fold dilutions were carried out from the 1:10 protein mixture at 4.75X prior to mixing with phase separation assay buffer. For reactions in supplementary figure 2 B and C, proteins (NUP98::KDM5A or NUP98::KDM5A W1625A) at a concentration of 750 nM were mixed with 0 or 2 µg chicken erythrocyte polynucleosomes in phase separation assay buffer. Similarly, *in vitro* FRAP experiments (Supplementary Figure 2E) were carried out with 1 µM NUP98::KDM5A mixed with 0 or 2 µg chicken erythrocyte polynucleosomes. Reactions with increasing concentrations of AF647-labeled H3K4me3 or H3K4me0 nucleosomal arrays (0-100 nM) (Figure 2 D-F and Supplementary Figure 2 F-H) were prepared by mixing varying concentrations of protein (0-800 nM of NUP98::KDM5A or NUP98::KDM5A W1625A) in phase separation assay buffer.

### Nucleosome assembly and *in vitro* phase separation assays: Microscopy

All images for in vitro phase separation assays in this study were acquired at the Center for Advanced Light Microscopy at UCSF. Confocal microscopy images were acquired using a CREST X-Light V2 large field of view spinning disk confocal unit, 100 X 1.40 NA oil objective, and Prime 95B 25mm sCMOS camera. Photobleaching was carried out with an Opti-Microscan FRAP unit using a 405nm laser line at 10% of the maximum laser intensity. Confocal images of *in vitro* reconstitutions are located in folder Figure2_ConfocalImaging_inVitro in the Dryad repository.

### Cell FRAP

Confocal live imaging coupled with fluorescence recovery after photobleaching (FRAP) of U2OS cells expressing mEGFP-NUP98::KDM5A and NUP98::KDM5A-mEGFP was performed in an imaging chamber maintained at 37 °C and 5% CO₂, mounted on an Andor Borealis CSU-W1 Spinning Disk Confocal microscope. We used a Plan Apo VC 100x/1.4 Oil objective. Imaging was carried out in a single channel using a 488 nm laser at 40% power (GFP channel, 200 ms exposure) with 1 × 1 binning. Images were acquired using a Zyla sCMOS camera (16-bit bit depth). For each field of view, time-lapse movies of 2048 × 2048 pixel images (pixel size 0.065 µm) were collected at a temporal resolution of approximately 220 ms per frame for over 30 s. Photobleaching was performed using a 405 nm laser (60 mW) with a 200 ms exposure time. Bleaching was applied approximately 3 s after the start of acquisition (∼10 frames) using four sequential laser pulses targeting a rectangular region of interest (∼150 × 50 pixels). Raw images used for Figure 2 - Figure Supplement 2 - A can be found in Figure2_ConfocalImaging_inVitro\Figure2-FigureSupplement2TIFF\A in the Dryad repository.

### Image analysis

Image analysis of condensates formed *in vitro* was carried out using FIJI (Version 1.54p) (Schindelin et al. 2012). Mean pixel intensities for protein condensates were calculated in FIJI from .tiff files collected under identical microscopy settings. Condensates were picked using an automatic threshold (3%) and the fraction of signal in condensates was determined by dividing the mean fluorescent protein intensity within condensates relative to the intensity in the entire image.

FRAP recovery curves for the photobleached region of interest (ROI) in both *in vitro* and in-cell experiments were generated by subtracting background fluorescence from the measured recovery signal and normalizing the data to the pre-bleach intensity. The FRAP recovery curves were fitted by non-linear regression using a one-phase exponential association model (GraphPad Prism 9.5.1, GraphPad Software, La Jolla, CA). In this model, the mobile fraction (M_F_) corresponds to the plateau value, while the half-life (T_1/2_) represents the time required for the fluorescence intensity to reach 50% of the maximum of the plateau level.

### Transfection procedure for cell imaging experiments

Sample preparation was performed as follows: 15 000–25 000 cells were plated into each of the wells of a Cellvis 8-chambered coverglass system (#1.5 polymer coverslip; Product#: C8-1.5P) for confocal and SoRa imaging, or Lab-Tek II 8-chambered coverglass system (#1.5 borosilicate coverglass; 155409) for Nikon-NSPARC live-imaging. Chambers were pre-treated with Poly-L-lysine solution (Sigma-Aldrich; P4707). Cells were plated in 200 µL culture medium and incubated for 18–24 h to allow attachment. The following day, 150–200 ng of plasmid DNA were transfected using the jetOPTIMUS DNA Transfection Kit (Item No. 101000025) according to the manufacturer’s short protocol (https://www.polyplus-sartorius.com/products/jetoptimus), adjusted for a 200 µL media volume. Cells were incubated for 4–6 hours in the transfection medium, after which the transfection medium was replaced with fresh culture medium and the cells were allowed to recover overnight. The following day, cells were either prepared for live-cell imaging experiments or fixed and processed for immunostaining and FISH protocols.

### Live cell imaging conditions

For live imaging, cells transfected with the appropriate NUP98::KDM5A constructs were incubated for 1 hour in FluoroBrite DMEM medium (Gibco #A1896701) supplemented with GlutaMAX (Gibco #35050-061) and SPY650-DNA stain (Cytoskeleton Cat. #CY-SC501). Samples were then imaged at 37 °C and X% CO₂ in cell-culture imaging chambers mounted on the corresponding imaging microscope.

### Immunostaining

For immunostaining, cells transfected with the appropriate NUP98::KDM5A–mEGFP or NUP98::KDM5Amut–mEGFP constructs were fixed with 4% PFA (Electron Microscopy 32% Paraformaldehyde, aqueous solution; #50980494) at room temperature for 10 min, PBS washed, and incubated for 1.5 hours at 37 C with a primary rabbit monoclonal antibody against the H3K4me3 histone mark (Cell Signaling Technology; C42D8) diluted 1:200. After PBS washes, samples were incubated for 1 hour at room temperature with a goat anti-rabbit secondary polyclonal antibody conjugated with Alexa Fluor 546 (Thermo Fisher; A11035) diluted 1:1000. Then, cells were stained with DAPI and post-fixed for 10 min at room temperature with 2% paraformaldehyde and 0.1% glutaraldehyde (Electron Microscopy Sciences; Cat #16019). All antibody incubations were performed in an immunostaining that contains PBS, 0.2% Triton X-100 (Sigma; T9284) and 3% Bovine Serum Albumin (BSA; Jackson Immuno Research; Code: 001-000-162), as described in the protocol available on Figure1_NUP98KDM5A_H3K4me3_correlation\ImmunostainingProtocol in Dryad repository.

### Combined FISH and Immunostaining

For FISH combined with immunostaining, cells transfected with a NUP98::KDM5A-FLAG construct were fixed with 4% PFA for 10 minutes at room temperature, washed with PBS, and processed through a FISH protocol based on (Mateo et al., 2019). First, samples were gently permeabilized for 10 minutes with 0.5% Triton-X, followed by a 1-hour RNase A (Monarch; Catalog# T3018L) treatment at 37 C and a 5-minute deproteinization step in 0.1 M HCl. After PBS washing, samples were incubated at 42 C for 30 minutes in a pre-hybridization buffer composed of 50% formamide (Millipore; #344206), saline-sodium citrate buffer (SSC; Invitrogen; Ref AM9763), Tween 20 (Sigma; P9416). Next, samples were transferred into a hybridization buffer composed of 50% formamide, SSC, Tween 20, and dextrane sulfate solution (Millipore; S4030) containing two sets of FISH probes targeting contiguous loci (5 ng/µL each; details in FISH probe design section). Genomic DNA was denatured on a heat block at 93 C for 5 minutes, after which samples were incubated at 42 C overnight. The following day, formamide was removed through a series of SSC washes, and the samples were immunostained by incubating them for 1.5 hours at 37 C with DYKDDDDK (Flag) Tag recombinant rabbit monoclonal antibody conjugated with Alexa Fluor Plus 488 (Thermo Fisher; 8H8L17) diluted 1:200 in immunostaining buffer. We performed a post-hybridization post-fixation step using 2% paraformaldehyde and 0.1% glutaraldehyde for 10 minutes at room temperature. Finally, we hybridized the secondary pairs of FISH probes (100 nM each), tagged with Cy3 and Cy5 and corresponding to each contiguous set of primary probes, by incubating the samples in ethylene carbonate (Sigma Aldrich; E26258) buffer. Samples were then washed with SSC buffer and 30% formamide, and stained with DAPI. For detailed protocol, see Figure3_FISHProtocol in Dryad repository.

### FISH probe design

FISH probes were designed by first retrieving pairs of 10-kb contiguous genomic regions corresponding to portions of the loci of interest: HOXA cluster (chr7:27197026-27207050 and chr7:27186295-27196321), HOXB cluster (chr17:46653759-46663990 and chr17:46665068-46675304), HOXD cluster (chr2:176957019-176967205 and chr2:176968092-176978322), TNF (chr6:31529154-31539396 and chr6:31539895-31550008), MUC4 (this probe is the oligo CCCCTCTTCCTGTCACCGACACTTCCTCAGCATCCACAGG, a repetitive sequence distributed among 11 kb within MUC4) and MYC (chr8:128739417-128749599 and chr8:128749901-128760113). Genomic regions were selected according to their H3K4me3 density in HEK293 cells, as assessed using the public ChIP-seq dataset “HEK293 H3K4me3 Histone Modifications by ChIP-seq from ENCODE/University of Washington” (Sabo et al., 2004). Genomic region sequences were retrieved from UCSC Genome Browser on Human (GRCh37/hg19) (Perez et al., 2025) and each region downloaded as a FASTA file. We then used the bioinformatic tool OligoMinerApp (http://oligominerapp.org/) (Passaro et al., 2020) to generate approximately 100 probes per 10-kb region, using the following parameters: MIN length 37 bp; MAX length 42 bp; MIN GC 35 %; MAX GC 70 %; MIN Tm 42 C; MAX Tm 47 C; Prohibited sequences AAAAA,TTTTT,CCCCC,GGGGG; MIN space 0 bp; [Na+] 390 Mm; [Formamide] 50%. All other parameters were set to default. Finally, we incorporated short overhang sequences into the primary probes to enable hybridization with fluorophore-labeled secondary probes. The overhangs were designed to be complementary to the secondary probe sequences: the “GCTTTAAGGCCGGTCCTAGC-Cy5” and “Cy5-AGCTCACCTATTAGCGGCTAAGG” secondary probes (conjugated to Cy5) hybridize to the overhangs flanking one set of primary probes, whereas the “AGCTGATCGTGGCGTTGATG-Cy3” and “Cy3-TGGACGGTTCCAATCGGATC” secondary probes (conjugated to Cy3) hybridize to the overhangs flanking the second, contiguous set of primary probes. Primary probes were synthesized as 50 pmole oligo pools, and secondary probes were synthesized as 100 nmole modified DNA oligos labeled with Cy3 or Cy5. Both sets of probes were synthesized by IDT. Synthesized probe sequences are provided in the [gene]-IDT .xlsx files within each [gene] folder in folder Figure3_DNASequences\FISHProbes in Dryad repository.

### Live imaging with a regular confocal microscope

Confocal live imaging of U2OS cells expressing the NUP98::KDM5A-related constructs shown in Figure 1SA was performed in an imaging chamber maintained at 37 C and 5% CO_2_, mounted on an Andor Borealis CSU-W1 Spinning Disk microscope controlled by the open-source software Micromanager (D. Edelstein et al., 2014). We used a Plan Apo VC 100x/1.4 Oil objective. Imaging was carried out in two sequential channels using a 488 nm laser at 20% power (GFP channel, 200 ms exposure) and a 640 nm laser at 50% power (Cy5 channel, 500 or 800 ms exposure), with 1 x 1 binning. Images were acquired using an Andor Zyla sCMOS 4.2 camera (16-bit bit depth, rolling shutter, 200 MHz readout speed). For each field of view, 2048 × 2048 pixel images (0.065 µm pixel size) were collected as Z-stacks spanning 14 µm centered around the focal center, with a step size of 0.26 µm, resulting in a total of 54 slices. Raw images in Figure 1 - Figure Supplement 1 are in folder Figure1_Confocal_live_imaging in Dryad repository.

### Live imaging with the SoRa spinning disk confocal microscope

SoRa-enhanced resolution live imaging of U2OS cells expressing NUP98::KDM5A constructs tagged with mEGFP at either the N- or C-terminus was performed in an imaging chamber maintained at 37 C and 5% CO_2_, mounted on a Yokagawa CSU-W1 SoRA Spinning Disk Confocal Microscope controlled by the open-source software Micromanager. We used an HC PL APO 100x/1.47 OIL CORR TIRF objective. Imaging was carried out in two sequential channels using a 488nm laser at 1.5 V (30% power) (GFP channel, 200 ms exposure) and a 640 nm laser at 1.5 V (30% power) (Cy5 channel, 200 ms exposure), with 1 x 1 binning. Images were acquired using a Prime BSI (PCIe) sCMOS camera (16-bit bit depth, open shutter) coupled with SoRa optics. For each field of view, 2300 × 2100 pixel images (0.0262 µm pixel size) were collected as Z-stacks spanning 8 µm centered around the focal center, with a step size of 0.26 µm, resulting in a total of 31 slices. Raw images used for Figure 2 - Figure Supplement 2B and C are stored in folder Figure2_SoRa_live_imaging in Dryad repository.

### Live imaging with the NSPARC scanning confocal microscope

NSPARC-enhanced resolution live imaging of U2OS cells expressing the NUP98::KDM5A– related constructs was performed in an imaging chamber maintained at 37 C and 5% CO_2_, mounted on a Nikon Ti2-E scanning laser microscope equipped with the Nikon Spatial Array Confocal (NSPARC) detector. We used an oil objective Plan APO D 60 x oil OFN25 DIC N2-1.42 NA. Imaging was performed sequentially in two emission channels: 524 nm (EGFP channel) or 593 nm (mCherry channel), and 677 nm (Cy5 channel), each at a laser power setting of 3.0. Laser scanning was performed in bidirectional mode with an integration setting of 2 and a pixel dwell time of 0.3 µs, and fluorescence emission was collected using a quad-band emission filter. The virtual pinhole size was set to 1.0, yielding an optical resolution of 0.19 µm. Imaging was performed with a zoom factor of 3.9, resulting in a pixel size of 0.073 µm (Nyquist sampling: 0.083 µm), and a scan area of 1024 × 1024 pixels (76 µm × 76 µm). NSPARC super-resolution mode was enabled, and Z-stacks spanning 6 µm were acquired with 13 slices at 0.5 µm step size. The frame acquisition time was 2.3 s per slice, resulting in a total stack acquisition time of approximately 30 s. Raw images for Figure 1 (A, B, and C), and Figure 1 - Figure Supplement 2A and B are stored in folder Figure1_NSPARC_live_imaging in Dryad repository.

### NSPARC imaging of immunostaining samples

NSPARC-enhanced resolution imaging of U2OS cells expressing NUP98::KDM5A-mEGFP or NUP98::KDM5Amut-mEGFP and immunostained for H3K4me3 was performed using an NSPARC scanning laser microscope. We used an oil objective Plan APO D 60 x oil OFN25 DIC N2-1.42 NA. Imaging was performed sequentially in three emission channels: 452 nm (DAPI channel, laser power 1.0), 524 nm (EGFP channel, laser power 4.0), and 593 nm (AF546 channel, laser power 2.0). Laser scanning was performed in bidirectional mode with an integration setting of 4 and a pixel dwell time of 0.3 µs, and fluorescence emission was collected using single emission filters. The virtual pinhole size was set to 1.0, yielding an optical resolution of 0.19 µm. Imaging was performed with a zoom factor of 3.9, resulting in a pixel size of 0.073 µm (Nyquist sampling: 0.083 µm), and a scan area of 1024 × 1024 pixels (76 µm × 76 µm). NSPARC super-resolution mode was enabled, and Z-stacks spanning 4 µm were acquired with 9 slices at 0.5 µm step size. Raw images for Figure 1 D, E and F can be found in folder Figure1_NUP98KDM5A_H3K4me3_correlation in Dryad repository.

### NSPARC imaging of combined FISH and immunofluorescence samples

NSPARC-enhanced resolution imaging of HEK293FT cells immunostained for NUP98::KDM5A– FLAG and labeled by FISH at specific genomic loci was performed using an NSPARC scanning laser microscope. We used an oil objective Plan APO D 60 x oil OFN25 DIC N2-1.42 NA. Imaging was performed sequentially in four emission channels: 452 nm (DAPI, laser power 3.0), 524 nm (AF488, laser power 3.0), 593 nm (Cy3, laser power 4.0), 699 nm (Cy5, laser power 4.0). Laser scanning was performed in unidirectional mode with an integration setting of 2 and a pixel dwell time of 0.4 µs, and fluorescence emission was collected using single emission filters. The virtual pinhole size was set to 1.0, yielding an optical resolution of 0.19 µm. Imaging was performed with a zoom factor of 2, resulting in a pixel size of 0.073 µm (Nyquist sampling: 0.083 µm), and a scan area of 2048 × 2048 pixels (150 µm × 150 µm). NSPARC super-resolution mode was enabled, and Z-stacks spanning 10 µm were acquired with 53 slices at 0.195 µm step size (Nyquist sampling on Z). Raw images for Figure 3B, C and D and Figure 3 - Figure Supplement 1A and B can be found under [gene] folders in Figure3_NSPARC_FISH_Imaging in Dryad repository.

### Calibration between mEGFP and mCherry signals

For mEGFP-to-mCherry calibration, we generated a DNA construct expressing the C-terminal domain of KDM5A (amino acids 1486–1690), which contains the PHD3 chromatin-binding domain and does not form condensates, tagged with mEGFP at the N-terminus and mCherry at the C-terminus. Transient transfection was performed in U2OS cells, taking advantage of the broad range of expression levels produced by transient expression. Transfected cells were imaged using the same protocol and settings employed for NSPARC live-cell imaging. Raw images for Figure 1 - Figure Supplement 2C are located in Figure1_mEGFP_mCherry_calibration.

Cells were segmented using Cellpose 2.0 (Pachitariu & Stringer, 2022), and mean fluorescence intensities in the mEGFP and mCherry channels were measured within the segmented nuclear masks across a range of expression levels using the script Figure1_Code\NSPARC-mEGFPmCherry-Calibration.ipynb. Output files with the suffix _cells and _parameters were saved in folder Figure1_mEGFP_mCherry_calibration\analysis_20241011-NSPARC_U2OSlive-mEGFP-KDM5A-mCherry. Then, we used the script Figure1_Code\Figure1A.ipynb to calculate a regression line relating mEGFP and mCherry intensity values acquired under NSPARC live imaging settings, and generated Figure 1 - Figure Supplement 2C. Paths correspond to the Dryad repository.

### EGFP intensity to concentration calibration

Sample preparation for intensity-to-concentration calibration was performed as follows: Sonicated TetraSpeck beads of 0.1 µm diameter (Invitrogen; Catalog #T7279) were diluted 1:1000 in PBS. Purified EGFP (Recombinant Purified Protein; ChromoTek; Cat. EGFP) was then diluted into the PBS–bead solution to final concentrations of 0 nM, 25 nM, 50 nM, 100 nM, 200 nM, 400 nM, 500 nM for SoRa analysis; and 200 nM, 100 nM, 50 nM for NSPARC analysis. EGFP has nearly identical molecular brightness as mEGFP. Each EGFP dilution was plated into a Lab-Tek II 8-chambered coverglass system (#1.5 borosilicate coverglass; 155409) for imaging under the same settings used for SoRa live imaging or NSPARC live imaging. Raw SoRa EGFP calibration data used for Figure 2 - Figure Supplement 2D is stored in Figure2_SoRa_EGFP_concentration_calibration. Raw NSPARC EGFP calibration data used for Figure 1 - Figure Supplement 2D is in Figure1_NSPARC_EGFP_concentration_calibration.

For each concentration of SoRa EGFP concentration to intensity analysis, we recorded one biological replicate and four technical replicates. Each replicate was recorded as 2300 x 2100 pixel images over a Z-stack of 41 slices with a step size of 0.5 µm, spanning a total of 20 µm, around the bottom of the chamber, as shown by the presence of TetraSpeck beads. Image analysis was performed in a Python-based Jupyter Notebook script Figure2_Code\SoRa-EGFPCalibration.ipynb. For each image in the Z-stack, a 1000 × 1000 pixel region centered within the field of view was extracted. The mean fluorescence intensity was calculated for each z-slice, and the 20th slice—corresponding approximately to the bottom of the chamber, as indicated by the presence of Tetraspeck beads—was selected. Fluorescence intensity was then quantified as the mean intensity of the 1000 × 1000 pixel region in the z-slice located 8 slices (approximately 4 µm) above this reference plane, ensuring measurement within the EGFP solution. Background subtraction was performed by subtracting the mean intensity of a 1000 × 1000 pixel region from a z-slice located 8 planes (approximately 4 µm) below the bottom of the chamber. We read output files, and the script Figure1_Code\FigureS1.ipynb to generate Figure 2 - Figure Supplement 2D.

As for NSPARC EGFP concentration to intensity analysis, we recorded two biological replicates and three technical replicates for all concentrations. Each replicate was recorded as 1024 x 1024 pixel images over a Z-stack of 63 slices with a step size of 0.195 µm, spanning a total of 12 µm, around the bottom of the chamber, as shown by the presence of TetraSpeck beads. Image analysis was performed in a Python based Jupyter notebook script Figure1_Code\NSPARC-EGFPCalibration.ipynb. For each image in the Z-stack, a 600 × 600 pixel region centered within the field of view was extracted. The mean fluorescence intensity was calculated for each z-slice, and the slice with the highest mean intensity—corresponding approximately to the bottom of the chamber as indicated by the presence of tetraspeck beads— was identified. Fluorescence intensity was then quantified as the mean intensity of the 600 × 600 pixel region in the z-slice located 20 slices (approximately 4 µm) above this reference plane, ensuring measurement within the EGFP solution sample. Because regions outside the sample yield zero intensity values under NSPARC imaging conditions, background subtraction was not required. An output file was generated and stored in Figure1_NSPARC_EGFP_concentration_calibration\20241202-NSPARC-EGFP_NSPARCcalibration_output.txt.

We used the output data to generate Figure 1 - Figure Supplement 2D through the script Figure1_Code\Figure1A.ipynb. Paths correspond to the Dryad repository.

### Image analysis: U2OS NSPARC live imaging

Raw image files were read and analyzed using custom scripts in a Python-based Jupyter Notebook script Figure1_Code\NSPARC-CondensatesQuantification.

For each dataset, the central z-slice was selected manually, and a single slice per dataset was used for downstream analysis. Nuclei were segmented using Cellpose 2.0 on the DAPI channel (Cell pose model: cyto), and nuclear masks were generated. Condensates were segmented from the corresponding fluorescence channel using a Difference of Gaussians (DoG) filter (low sigma: 1.5 pixels; high sigma: 2 pixels), followed by thresholding in DoG space (gaussian threshold for mEGFP: 1e-05, gaussian threshold for mCherry: 4e-06). Segmented condensates smaller than 4 pixels were excluded from further analysis. Remaining condensates were assigned to individual nuclear masks, and quantitative statistics were computed for both nuclei and condensates. Nuclear measurements included x and y coordinates, total nuclear fluorescence intensity (raw_exp), nuclear area (cell_area), background intensity (background), nucleoplasmic intensity (nuc_intensity), and nucleoplasmic area (nuc_area), where the nucleoplasm was defined as the portion of the nucleus excluding condensates after a 3-pixel erosion. Condensate measurements included x and y coordinates, integrated intensity (raw_mass), size (size), and eccentricity (ecc). The resulting measurements were exported for downstream analysis into separate output .txt files with the suffixes _cells, _condensates, and _parameters, which contain details of the analysis. All files were saved within the Analysis folder of each dataset. Images of the segmented nuclei were saved in the Images subfolder. Figure 1C and Figure 1 - Figure Supplement 2B were generated with the output files and the script Figure1_Code\Figure1A.ipynb. Paths correspond to the Dryad repository.

### Image analysis: U2OS SoRa live imaging

Segmentation and quantification of SoRa-imaged condensates were performed as described for the NSPARC datasets, with modifications, using script Figure2_Code\SoRa-CondensatesQuantification. Nuclei were segmented using Cellpose 2.0 on the Cy5 channel (Cellpose model: cyto). Condensates were segmented using a Difference of Gaussians filter (low sigma: 1.5 pixels; high sigma: 4.5 pixels), followed by thresholding in DoG space (Gaussian threshold for mEGFP: 1e-04). Segmented condensates smaller than 4 pixels were excluded. The nucleoplasm was defined as the portion of the nucleus excluding condensates after a 5-pixel erosion. Nuclear and condensate measurements were identical to those used for U2OS NSPARC live imaging analysis. The resulting measurements were exported for downstream analysis into separate output .txt files with the suffixes _cells, _condensates, and _parameters, which contain details of the analysis. All files were saved within the Analysis folder of each dataset. Images of the segmented nuclei were saved in the Images subfolder. We read output files and generated Figure 2 - Figure Supplement 2C with the script Figure1_Code\FigureS1.ipynb. Paths correspond to the Dryad repository.

### Image analysis: Colocalization analysis between NUP98::KDM5A and H3K4me3

Raw image files were read and analyzed using the Python-based Jupyter Notebook script Figure1_Code\NSPARC-NK-H3K4me3-Correlation.ipynb.

For NSPARC microscopy of U2OS cells expressing NUP98::KDM5A-mEGFP or NUP98::KDM5Amut-mEGFP and immunostained for H3K4me3, images were analyzed using Python-based Jupyter Notebook scripts. After manual selection of the central Z-slice, channel alignment between NUP98::KDM5A and H3K4me3 was corrected using phase cross-correlation. This method leverages the cross-power spectrum and inverse Fourier transform to estimate inter-channel offsets with subpixel accuracy. Following channel alignment, nuclear masks were generated using Cellpose 2.0 (model: cyto) applied to the H3K4me3 channel after Gaussian smoothing. Fluorescence intensities from the H3K4me3 and NUP98–KDM5A channels were extracted from each individually segmented nucleus, flattened into one-dimensional vectors, and zero-valued pixels were discarded. Pearson’s correlation coefficient was then computed to quantify pixel-level linear association between the two signals. For visualization, two-dimensional histograms of pixel intensity values were generated (x-axis: mEGFP-NUP98::KDM5A intensity; y-axis: H3K4me3–AF546 intensity) and rendered using logarithmic color normalization to emphasize differences in pixel-density distributions. The highest density typically appeared near the origin, corresponding to the background signal. We generated output files named megfp_546_with_iou.txt and megfpmut_546_with_iou.txt, which contain tabulated measurements including the Pearson correlation coefficient between channels (Pearson_r), mean nuclear fluorescence intensity (Expression), and the intersection-over-union (IoU) coefficient between the two channels. Two-dimensional histograms were saved in folders named 2D_histograms, located in the same directory as the corresponding output files. These files are located in Figure1_NUP98KDM5A_H3K4me3_correlation\NSPARC_Nup98_KDM5A_Correlation_Analysis. Output files were analyzed with Python-Jupyter notebook script Figure1_Code\Figure1B.ipynb to generate Figure 1F. Paths correspond to the Dryad repository.

### Image analysis: FISH analysis

NSPARC images of immunostaining and DNA FISH samples were analyzed. Images were acquired at 2048 × 2048 pixels, corresponding to a field of view of 150 µm × 150 µm, as Z-stacks spanning 10 µm with 53 slices. Each dataset contained four channels: DAPI (nuclear marker), AF488 (NUP98::KDM5A condensates), and Cy3 and Cy5 channels corresponding to contiguous FISH probes labeling specific genomic loci. For selected datasets, a minimum 3D filter was applied with Fiji/ImageJ to enhance nuclear contrast prior to segmentation (kernel size: x = 3.0, y = 3.0, z = 1.0). These datasets are denoted by “m3d” in the segmentation mask file title. Three-dimensional nuclear segmentation was then performed using Cellpose version 4.0.3 in 3D mode with the CPSAM mode (Pachitariu et al., 2025). Masks were saved as .tiff files with the suffix _cp_masks.

To detect FISH-labeled genomic loci, we used the Python-based package Big-FISH (Imbert et al., 2021) implemented in Jupyter Notebook scripts. Big-FISH was applied to the Cy3 and Cy5 channels to localize contiguous genomic loci in three dimensions, extracting their x, y, and z coordinates from the full Z-stacks for downstream spatial analyses. In short, the Cy3 and Cy5 channels were first thresholded at 2 au, followed by three-dimensional Gaussian filtering (σ = 1 pixel in x, y, and z) for denoising. Local maxima detection was then applied to identify Gaussian-like intensity spots corresponding to FISH signals. A maximum was defined as a pixel whose value remained unchanged before and after application of the local maxima filter, with a minimum distance of 4 pixels enforced between detected maxima. Detected maxima were further filtered by intensity to retain only the brightest peaks (Cy3 ≥ 3 au; Cy5 ≥ 5 au). To identify legitimate probe pairs hybridized to contiguous genomic loci, only Cy3 and Cy5 peaks located within 300 nm of each other were retained, based on voxel dimensions of 0.073 µm × 0.073 µm × 0.195 µm (x, y, z). For each valid probe pair, the midpoint between the Cy3 and Cy5 coordinates in three-dimensional space was defined as the final x, y and z positions of the genomic locus, denoted as cy5cy3_x, cy5cy3_y, cy5cy3_z. Output files, named with the suffix _fish and _fish_supplement were generated for downstream analyses. This part of the analysis was performed using Python-based Jupyter Notebook script Figure3_Code\FISHPairs.ipynb.

To study the association between NUP98::KDM5A and FISH-labeled genomic loci, we read nuclear segmentation masks generated by Cellpose (files with the suffix _cp_masks) and corresponding locus coordinate files (files with the suffix _fish) containing the three-dimensional positions of genomic loci. Nuclear masks were applied to the AF488 channel, which shows NUP98::KDM5A condensates, and the mean NUP98::KDM5A fluorescence intensity within each nucleus was measured (Mean NUP98::KDM5A intensity in cells). FISH spots were then assigned to their corresponding nuclei based on 3D spatial overlap within the cellpose masks, enabling quantification of NUP98–KDM5A signal in relation to the labeled genomic loci.

For quantification of NUP98::KDM5A intensity around individual genomic loci, ellipsoidal volumes were modeled around each detected FISH spot. Because of anisotropic resolution of optical imaging, a spherical neighborhood in physical space corresponds to a flattened ellipsoid along the Z axis in voxel space. Voxel dimensions were 0.073 µm in X, 0.073 µm in Y, and 0.195 µm in Z, and ellipsoid dimensions were adjusted accordingly to accurately sample the local NUP98::KDM5A signal surrounding each locus. For each locus, we computed the mean, maximum, and 20th percentile value of NUP98::KDM5A fluorescence intensities within ellipsoids of radii 100, 200, 300, 400, 800, and 1000 nm. These measurements were saved in output .txt files with the suffix _radial and _radial_supplement. Images of individual FISH spots and surrounding regions were saved in folders named after the corresponding dataset, located alongside the _radial files. This analysis was performed using the script Figure3_Code\RadialAnalysis.ipynb. Figure 3B was generated using a modified version of the script Figure3_Code\RadialAnalysis-Figure3.ipynb.

Finally, the intensity around locus, shown in Figures 3C and D, was calculated as the maximum pixel intensity within a 200 nm ellipsoid—representing the local condensate signal—minus the 20th percentile intensity within an 800 nm ellipsoid, which served as an estimate of the local background fluorescence. As a control, the same analysis was repeated using the unrelated DAPI channel instead of the AF488 channel (Figure 3SA and Figure S3B). As expected, the resulting measurements did not follow the same trends observed for AF488. Plots in Figure 3 and Figure 3 - Figure Supplement 1 were generated using the script Figure3_Code\Figure3.ipynb. Paths correspond to the Dryad repository.

### Single cell RNA seq analysis: Datasets

Single-cell RNA-seq data were analyzed from published datasets comprising pediatric leukemias with NUP98 onco-fusions (Umeda et al., 2025, GEO GSE287716) and bone marrow cells from healthy donors (H. Li et al., 2025, GEO: GSE189161). Two patients with NUP98::KDM5A fusion (SJALL040121 and SJAML061263) were considered, and healthy donor published dataset (H. Li et al., 2025), matched for age (pediatric patients aged 2, 4, 10, and 12 years). Annotated cell-state were selected based on the dominant cell-types in each patient. The matched comparator for SJAML061263 (referred to as patient 1 here) was composed of erythroid/megakaryocytic progenitors together with early stem/progenitor compartments (i.e.,’Mk/E-MPP’,’Mk-Prog’,’E-Prog-1’,’E-Prog-2’,’E-Prog-3,’My-MPP’,’MPP-1’,’MPP-2’,’MPP-3’, donor age: 2 and 4). While, patient SJALL040121 (referred to as “patient 2”, here) was composed of lymphoid progenitors and early stem/progenitor compartments (i.e., ’Ly-Prog-1’,’Ly-Prog-2’,’Ly-Prog-3’,’Ly-Prog-4’,’Ly-Prog-5’, ’MPP-1’,’MPP-2’,’MPP-3’ with donor age: 10 and 12 years). These matched subsets were used for all the downstream analysis. Relevant data was stored in the file Figure4_CombinedData_Tables_Fig4_Fig4S1_Fig1S3 for further analysis.

### Single cell RNA seq analysis: Data processing

Standard single-cell quality control procedures were applied, cells with high mitochondrial transcripts were excluded by (>20%), cells with low gene detection (i.e., < 200 detected genes per cell) and genes detected in at least 3 cells were included. Filtered counts were normalized for sequencing depth to a library size (10,000 counts per cell) and log transformed. No additional correction was applied.

### Single cell RNA seq analysis: Estimation of NUP98::KDM5A expression in patient cells

To quantify the relative abundance of fusion partner transcripts we computed KDM5A to NUP98 ratios in each group. This was computed as the ratio of mean expression of KDM5A to the mean expression of NUP98 on library-size normalized counts. Because transcripts are quantified by poly-A capture, transcript identity is determined by the 3′ end. As a result, the NUP98::KDM5A fusion transcript is classified as KDM5A by our analysis pipeline. This shifts the KDM5A-to-NUP98 ratio from approximately 1 in healthy donors, who harbor two NUP98 and two KDM5A alleles, to a ratio of nearly 2 in Patient 1 (AMKL) and Patient 2 (ALL), both of which carry one NUP98 allele, one KDM5A allele, and one NUP98::KDM5A fusion allele that we quantified as a second KDM5A allele (Figure 1 - Figure Supplement 3A). We also computed per-cell ratios for cells with detectable NUP98 abundance (>0) and assessed statistical significance using the Wilcoxon test for paired groups, evaluating differences in relative abundance between healthy and patient donors as well as within healthy and patient individuals (Figure 1 - Figure Supplement 3B, Figure4_CombinedData_Tables_Fig4_Fig4S1_Fig1S3; Data Table 3). Because the KDM5A-to-NUP98 ratio in patients is close to 2, this suggests that NUP98::KDM5A expression is approximately half that of wild-type NUP98. This inference is supported by the fact that NUP98::KDM5A expression is driven by the NUP98 promoter. Given OpenCell measurements indicating that endogenous NUP98 expression in HEK293T cells is ∼360 nM (Cho et al., 2022), we infer that NUP98::KDM5A is expressed at ∼180 nM in this system. We observed condensate formation at comparable concentrations in both U2OS and HEK293FT cells.

### Single cell RNA seq analysis: Differential gene expression analysis

Differential gene expression was assessed by comparing each patient sample to its matched healthy reference subset using the Wald test (diffxpy in scanpy). Genes with FDR-adjusted p-value ≤ 0.05 and absolute fold-change (i.e., Log2 fold-change) ≥ 0.50 were defined as differentially expressed genes (DEG files stored at Figure4_CombinedData_Tables_Fig4_Fig4S1_Fig1S3; Data Table 1-2).

To account for the limited number of donors, pseudo-replication, and inter-donor variability, we evaluated additional metrics to assess concordance in direction between gene expression and methylation within donors. Specifically, we performed a) DEG analysis for each patient separately with healthy donors (n = 2), and b) comparisons to individual donors using (i) all cells and (ii) bootstrap-based sub-sampling of both healthy and patient cells. Gene-level log2FC values were consistent using pooled or individual donor references and were robust to bootstrap sampling. Correlations were moderate to high for patient-versus-donor comparisons (P1: 0.71– 0.99; P2: 0.74–0.92 with respective healthy controls), but lower between healthy donors (P1: 0.64; P2: 0.56). Discordant genes (|log2FC| ≥ 0.45) were infrequent (P1: 1.15%; P2: 6.7%) and further reduced using pooled donors (≤0.48%). Notably, key gene sets (e.g., HOX genes) remained concordant in direction across all comparisons.

### Single cell RNA seq analysis: H3K4me3 intensity quantification from CUT&RUN assays

Total H3K4me3 signal per gene in patients was quantified using the published CUT&RUN data from Umeda et al. 2025 (GEO: GSE287298). Gene annotations and transcription start site (TSS) coordinates were obtained from a GTF file (GRCh38.p14). For each gene, a defined window centered on the annotated TSS was used. H3K4me3 peak coordinates from BED files (broad peaks, 2 replicates) were intersected with this window, and gene-level H3K4me3 intensity was calculated from BigWig tracks as the sum of signal intensities across all overlapping peaks within the defined window.

Genes were defined as H3K4me3 marked genes, if detected within ± 1kbp of the TSS of a gene since H3K4me3 is promoter-proximal mark. Total H3K4me3 intensities for H3K4me3 marked genes was computed by integrating the intensities across all the peaks in the Transcription start site (TSS-centered windows). For the downstream analysis, H3K4me3 marked genes were considered.

### Single cell RNA seq analysis: Window-size selection

Total window size was varied from (±TSS centered symmetric) 1kbp -75kbp i.e. (2kbp to 150kb in total) for H3K4me3 intensities calculations, since some genes span H3K4me3 breadth up to 60-100kbp (Benayoun et al., 2014). Cut-off or saturating window size was determined as the window size that first crosses the saturating H3K4me3 intensity mark defined as 90% of maximal value. High H3K4me3 intensity threshold was arbitrarily defined as 60th percentile from the pooled H3K4me3 signal distribution across all the window sizes.

### Single cell RNA seq analysis: Correlation between Gene-expression and H3K4me3 methylation

Genes defined as H3K4me3 marked genes were used to evaluate the relationship between the gene-specific H3K4me3 intensity in patients (at ± 50kb window centered around TSS), and the relative change in its expression. Data was binned based on H3K4me3 intensity, mean and standard error of mean computed within each bin.

#### Mean estimates and proportion of cells with detected expression

For each individual donor, gene-level mean expression and the fraction of cells with detectable expression (i.e., non-zero counts for that gene) were computed across all cells. Then genes were grouped into bins based on the H3K4me3 intensity at 100kbp, and averaged across all the genes within that bin.

#### Relative change in expression in patients compared to healthy donors

For a control baseline, for each gene, we randomly sampled a locus on the same chromosome with the same window size, excluding the TSS-centered window and similarly computed the binned statistics (Data as in Figure 4B; Figure4_CombinedData_Tables_Fig4_Fig4S1_Fig1S3; Data Table 4). Next, we ranked all the H3K4me3 marked genes by their intensity (at ± 50kb with highest intensity ranked first, filtered by adjusted p-value from differential gene expression by FDR-adjusted p-value ≤ 0.05), with color representing the log-fold-change in the gene expression. Only protein-coding HOXA/HOXB genes are considered to compute statistics, while all the other genes in the ranked set are grouped under “other genes”. Data for these genes with H3K4me3 intensity in 2000 and 100 kbp window-size and corresponding change in gene expression (Data as in Figure 4C; Figure4_CombinedData_Tables_Fig4_Fig4S1_Fig1S3; Data Table 5 and 6 for each patient).

## Data and Code Availability

Supplemental data files are available in the Dryad repository “Preferential formation of NUP98-KDM5A condensates at specific H3K4me3-rich loci drives leukemogenic gene expression - Data and Code” (DOI: 10.5061/dryad.m63xsj4hb). Specific file paths are indicated throughout the Materials and Methods section of the manuscript.

## Author Contributions

AB - Conceptualization, Data curation, Software, Formal analysis, Investigation, Visualization, Methodology, Writing – original draft, Writing – review and editing, Project administration

JES - Conceptualization, Data curation, Software, Formal analysis, Investigation, Visualization, Methodology, Writing – original draft, Writing – review and editing, Project administration

NK - Data curation, Software, Formal analysis, Investigation, Visualization, Methodology, Writing – original draft, Writing – review and editing

AM - Data curation, Software, Formal analysis, Visualization, Investigation, Methodology

TW - Formal analysis, Investigation, Methodology

CM - Formal analysis, Investigation, Methodology, Writing – review and editing

HL – Resources, Supervision, Funding acquisition, Writing – review and editing

GJN – Resources, Supervision, Funding acquisition, Writing – review and editing

DGF – Conceptualization, Resources, Supervision, Funding acquisition, Methodology, Writing – original draft, Writing – review and editing, Project administration

BH - Conceptualization, Resources, Supervision, Funding acquisition, Methodology, Writing – original draft, Writing – review and editing, Project administration

## Acknowledgements

Data for this study were acquired at the Center of Advanced Light Microscopy at UCSF on a W1-CSU Confocal obtained using NIH S10 Shared Instrumentation grant (1S10OD017993-01A1), on CREST/C2 Confocal obtained using grants from the UCSF Program for Breakthrough Biomedical Research funded in part by the Sandler Foundation and the UCSF Research Resource Fund Award, and at the UCSF Innovation Core at the Weill Institute for Neurosciences on a Yokagawa CSU-W1 SoRA Spinning Disk Confocal Microscope.

Data for this study were acquired in the Nachury laboratory at UCSF using a Nikon Ti2-E scanning laser microscope equipped with a Nikon Spatial Array Confocal (NSPARC) detector. We thank Maxence Nachury and members of the Nachury lab for providing access to their Nikon NSPARC system.

We thank members of the Huang, Fujimori, and Li laboratories at UCSF, particularly Daniel Foyt, Nitya Kopparapu, Yiming Kuang, Catherine Taylor, and Aivy David, for helpful discussions and support. We also thank Alistair Boettiger and members of the Boettiger laboratory at Stanford University for guidance in establishing our FISH protocol, in particular Kavvya Gupta and Derek Le. We further acknowledge the faculty of the Cold Spring Harbor Laboratory course *“Quantitative Imaging: From Acquisition to Analysis”* for insightful advice on imaging analysis.

B.H. and G.N. are supported by the NIH 4D Nucleome Consortium (U01DK127421). J.E.S. is supported by NIH Institutional Research and Academic Career Development Award (IRACDA) K12GM081266 through the UCSF IRACDA Scholars Program. D.G.F. is supported by the Bowes Biomedical Research Award. N.K. and H. L. are supported by NIH (R01AG083524, R33AG078793). B.H. and H.L. are Biohub San Francisco Investigators.

---

**Figure 1 Figure Supplement 1.**
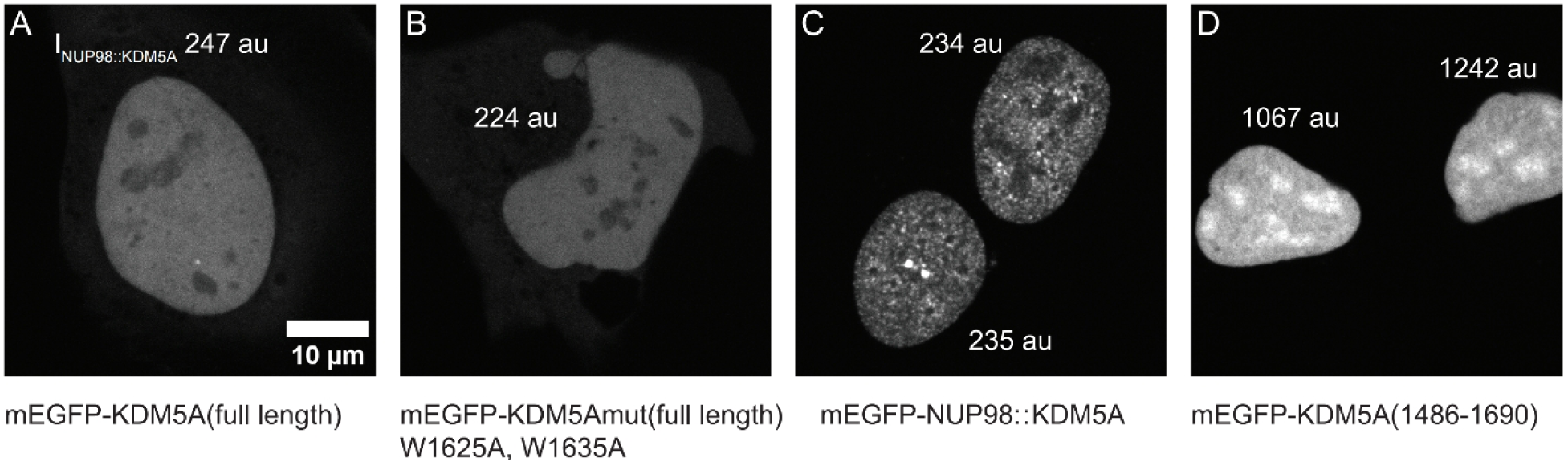
Live imaging of U2OS cell nuclei expressing NUP98::KDM5A-related constructs. **A.** Nucleus expressing full-length KDM5A construct fluorescently tagged on the N-terminus. **B.** Chromatin-binding-deficient version of the construct in A. **C.** Condensates formed by NUP98::KDM5A tagged with mEGFP on the N-terminus. **D.** Nuclei expressing only the chromatin-binding C-terminal fragment of the NUP98::KDM5A fusion, derived from KDM5A (1486–1690). A-C show cells with comparable expression levels, whereas D shows two cells with high expression but no chromatin-associated puncta formation. All images are a single z-slice acquired on a standard confocal microscope.

**Figure 1 Figure Supplement 2.**
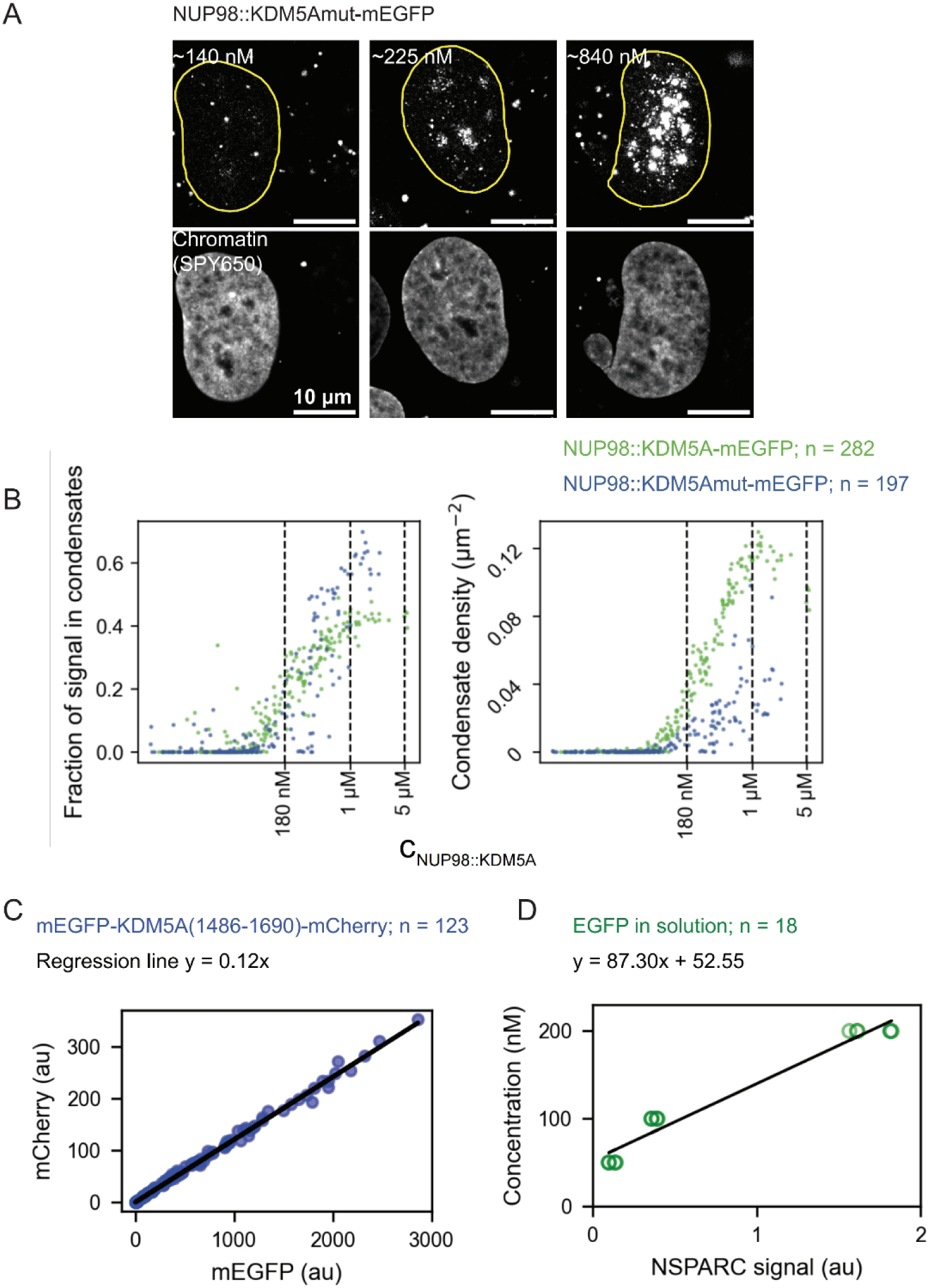
**NSPARC enhanced-resolution live imaging of NUP98::KDM5A condensates in U2OS cells**. **A.** Concentration-dependent formation of NUP98::KDM5Amut-mEGFP condensates, showing single z-slices from 3D stacks. **B.** Quantification of NUP98::KDM5A-mEGFP and NUP98::KDM5Amut-mEGFP condensate formation across expression levels for the fraction of signal in condensates and the number density of condensates. **C.** Calibration between mEGFP and mCherry fluorescence intensities under the NSPARC microscopy settings used for live-cell imaging (see Materials and Methods). **D.** Calibration between EGFP fluorescence intensity and EGFP molar concentration in solution under the NSPARC microscopy settings used for live-cell imaging (see Materials and Methods).

**Figure 1 Figure Supplement 3.**
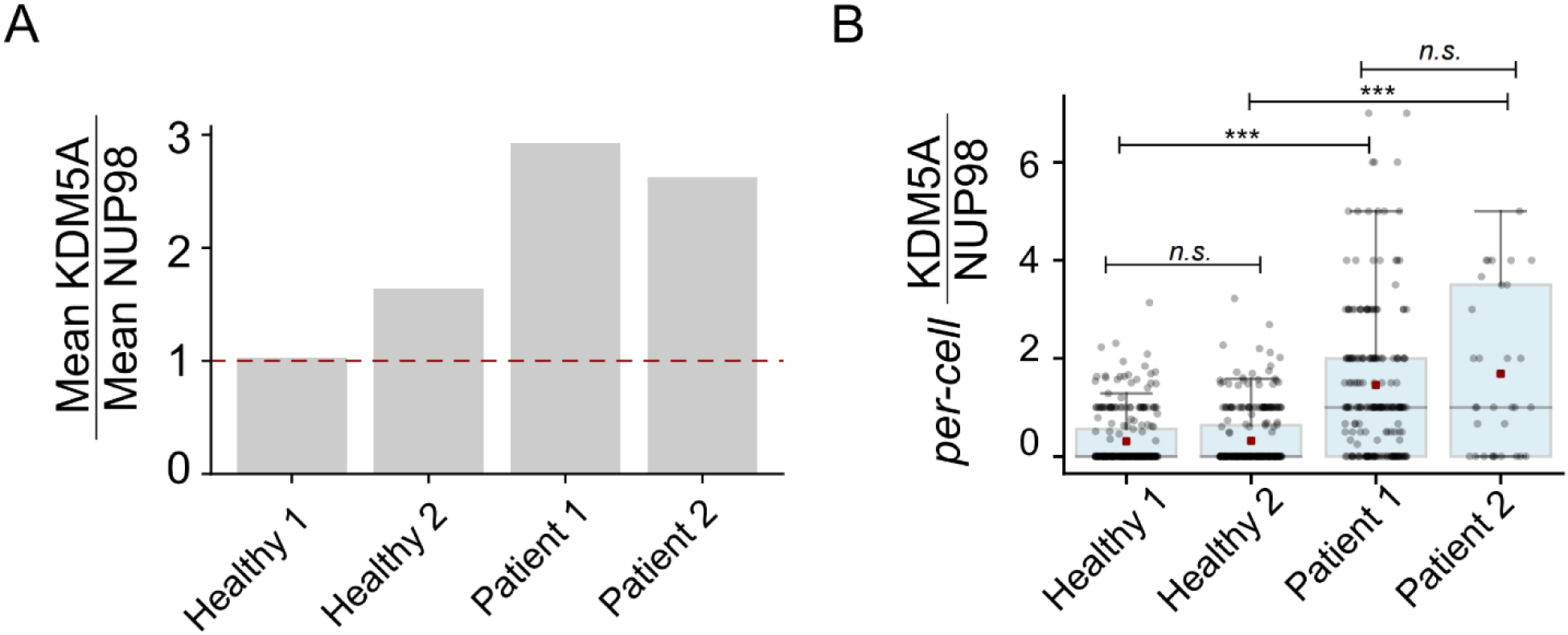
Estimation of the relative abundance of KDM5A to NUP98 in patient and healthy donors. **A.** Group level relative abundance of KDM5A: NUP98 transcripts quantified as the ratio of the mean expression across cells in each group. **B.** Distribution of per-cell KDM5A: NUP98 ratios for cells with detectable NUP98 expression (NUP98 > 0) across healthy and patient groups. Gray circles represent ratios from single-cells, and boxplots show distribution within each group. Red squares indicate the group mean, horizontal lines indicate medians, and whiskers extend to maximum points in 1.5 interquartile range (IQR). Wilcoxon rank sum test-ns: not significant, ***p< 0.001.

**Figure 2 Figure supplement 1.**
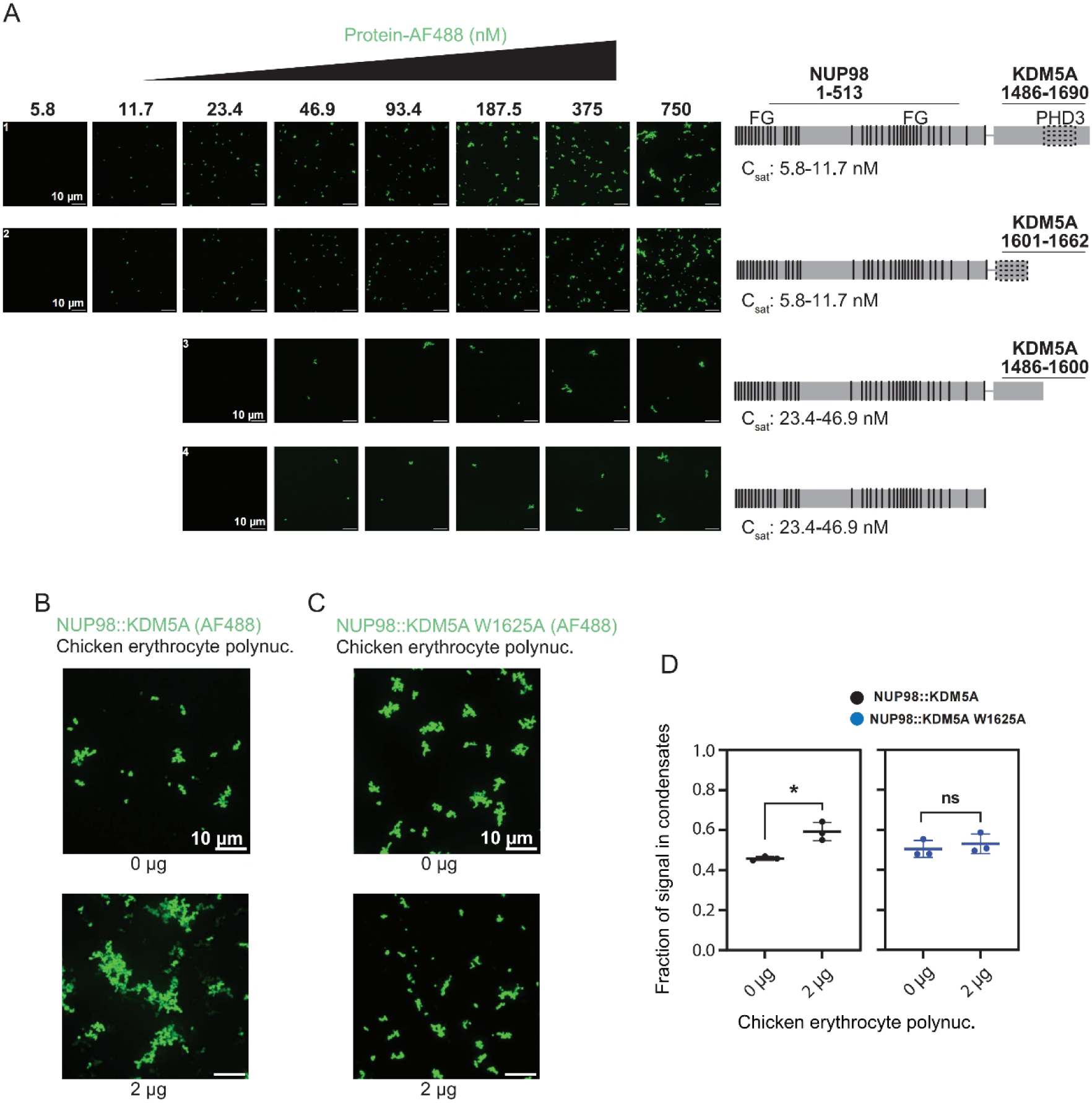
*In vitro* reconstitution of NUP98::KDM5A condensates. **A.** Spinning disk confocal images of condensates formed *in vitro* with increasing concentration of the following AF488 labeled constructs: NUP98 (1-513)-KDM5A (1486-1690), the full length oncofusion; NUP98 (1-513)-KDM5A (1601-1662), which contains only the PHD3 domain of the KDM5A fusion partner; NUP98 (1-513)-KDM5A (1486-1690), where the C-terminal truncation of the KDM5A fusion partner removes the PHD3 domain; NUP98 (1-513), the NUP98 segment alone. **B.** *In vitro* phase separation assay of AF488 labeled NUP98::KDM5A (750 nM) with chicken erythrocyte polynucleosomes (0 or 2 µg). **C.** *In vitro* phase separation assay of AF488 labeled NUP98::KDM5A W1625A (750 nM) with chicken erythrocyte polynucleosomes (0 or 2 µg). All images in **A**-**C** show maximum intensity projections of Z-stacks (scale bar = 10 µm; n = 3). **D.** Fraction of signal in condensates for reactions with 750 nM AF488 labeled NUP98::KDM5A with 0 or 2 µg chicken erythrocyte polynucleosomes (left panel) or 750 nM AF488 labeled NUP98::KDM5A W1625A with 0 or 2 µg chicken erythrocyte polynucleosomes (right panel). The fraction of signal in condensates corresponds to the fluorescence intensity measured inside condensates relative to the total image intensity, reflecting the proportion of molecules partitioned into condensates. A parametric unpaired t-test was carried out to compare the average for the fraction of signal in condensates (n=3; ns, P>0.05; *, P≤0.05; ** P≤0.01).

**Figure 2 Figure supplement 2.**
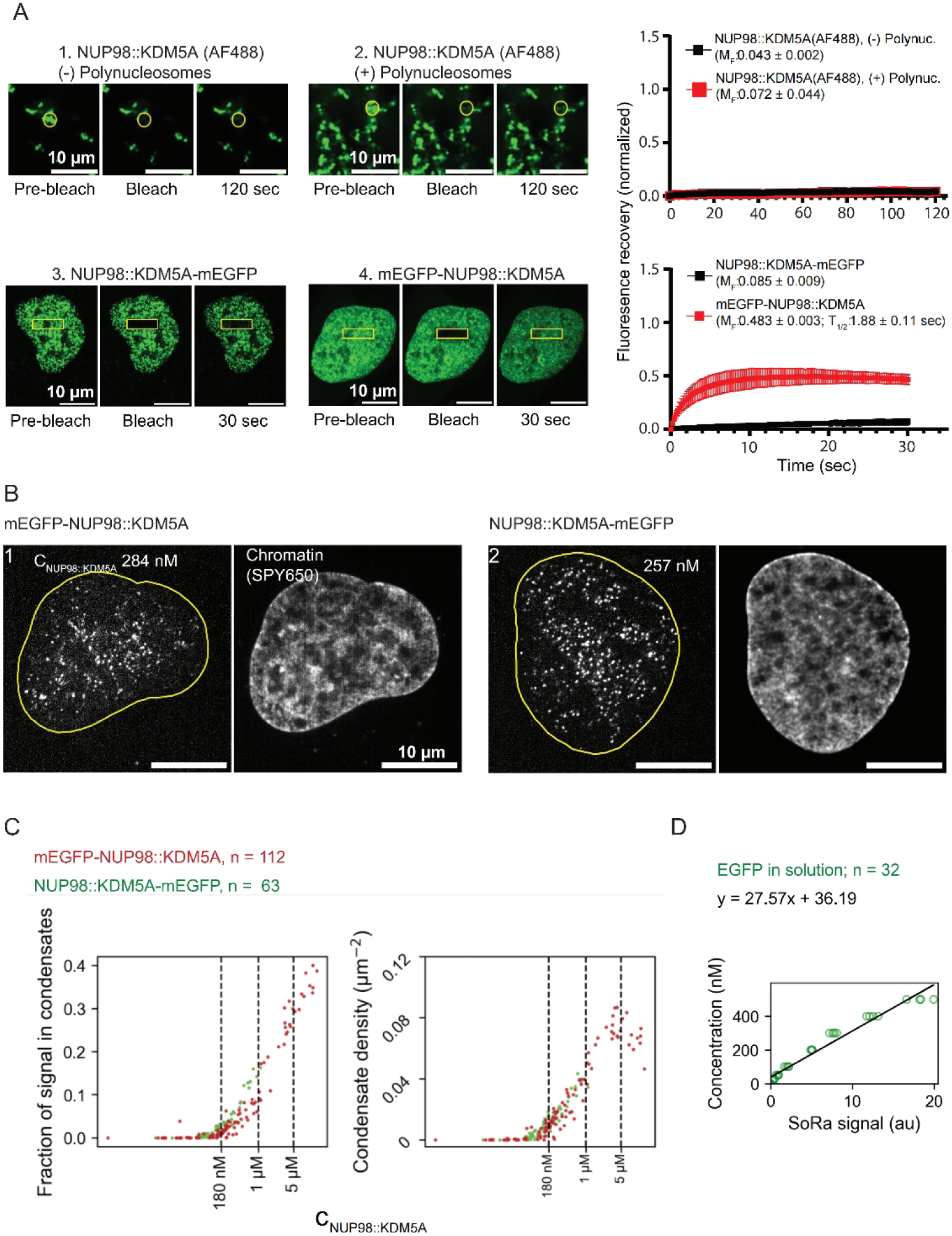
Dynamics of NUP98::KDM5A condensates. **A.** Representative images of the fluorescence recovery (inside yellow circle) of AF488 labeled NUP98::KDM5A (1 µM) mixed with 0 or 2 µg chicken erythrocyte polynucleosomes following photobleaching. Fluorescence recovery (inside yellow box) of U2OS cells expressing NUP98– KDM5A constructs tagged with mEGFP on either the C- or N-terminus after photobleaching. Normalized fluorescence recovery curves of photobleached regions (inside yellow marks) are plotted. Error bars are standard deviations of the means. **B.** SoRa-enhanced resolution live imaging of U2OS cells expressing NUP98::KDM5A constructs with an N- or C-terminal mEGFP tag. **C.** Quantification of mEGFP-NUP98::KDM5A and NUP98::KDM5A-mEGFP condensate formation across expression levels (C _NUP98::KDM5A_ - Log _10_) for the fraction of signal in condensates and condensate density (µm^-2^). **D.** Calibration between EGFP fluorescence intensity and EGFP molar concentration in solution under the SoRa microscopy settings used for live-cell imaging (see Materials and Methods).

**Figure 2 Figure supplement 3.**
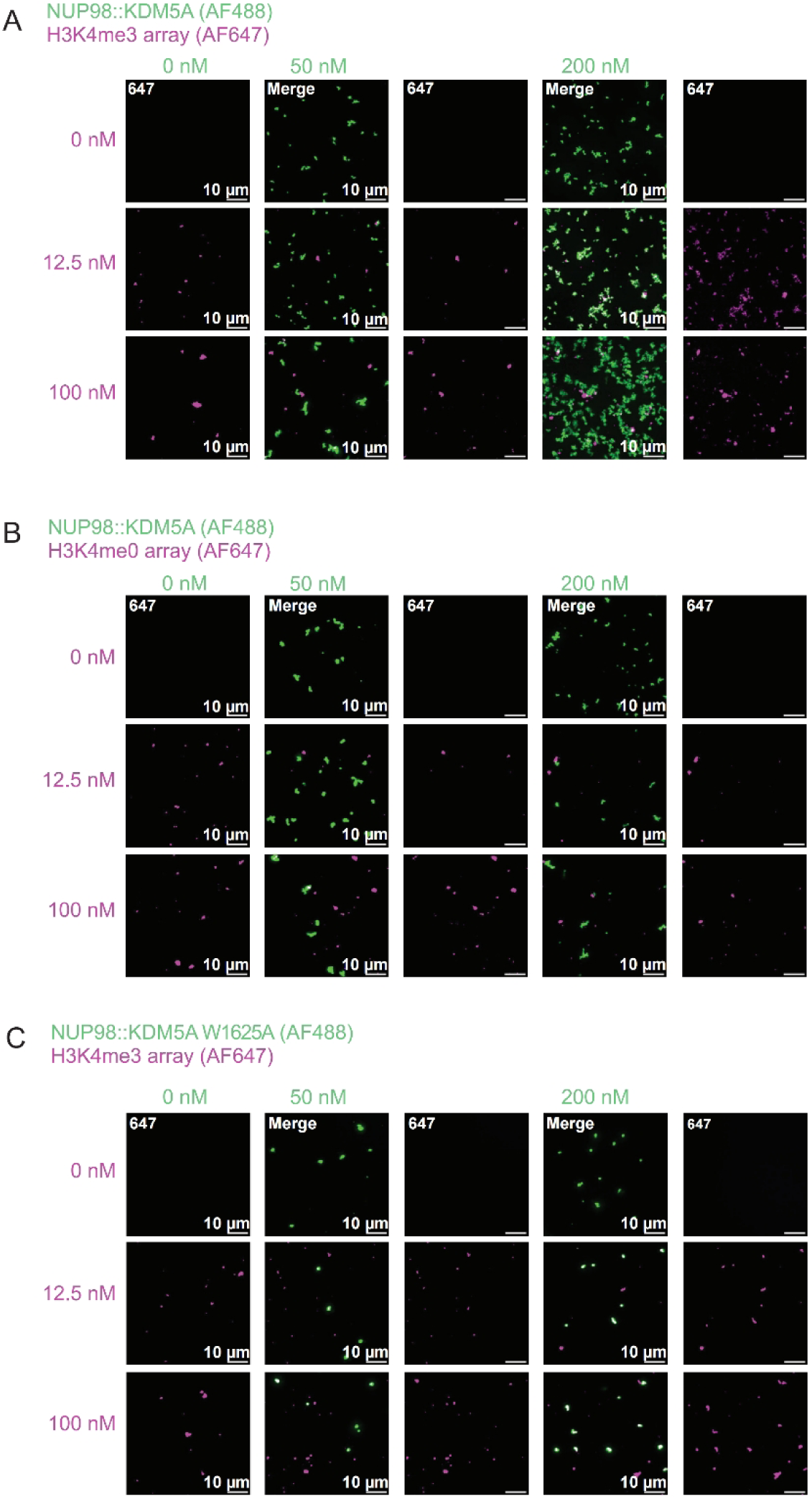
Representative images of *in vitro* NUP98::KDM5A and chromatin co-phase separation assays. **A.** AF488 labeled NUP98::KDM5A (0, 50, or 200 nM) with AF647 H3K4me3 nucleosomal arrays (0, 12.5, or 100 nM nucleosomes). **B.** AF488 labeled NUP98::KDM5A (0, 50, or 200 nM) with AF647 H3K4me0 nucleosomal arrays (0, 12.5, or 100 nM nucleosomes). **C.** AF488 labeled NUP98::KDM5A W1625A (0, 50, or 200 nM) with AF647 H3K4me3 nucleosomal arrays (0, 12.5, or 100 nM nucleosomes). Micrographs are presented as maximum intensity projections (scale bar = 10 µm; n = 3).

**Figure 3 Figure Supplement 1.**
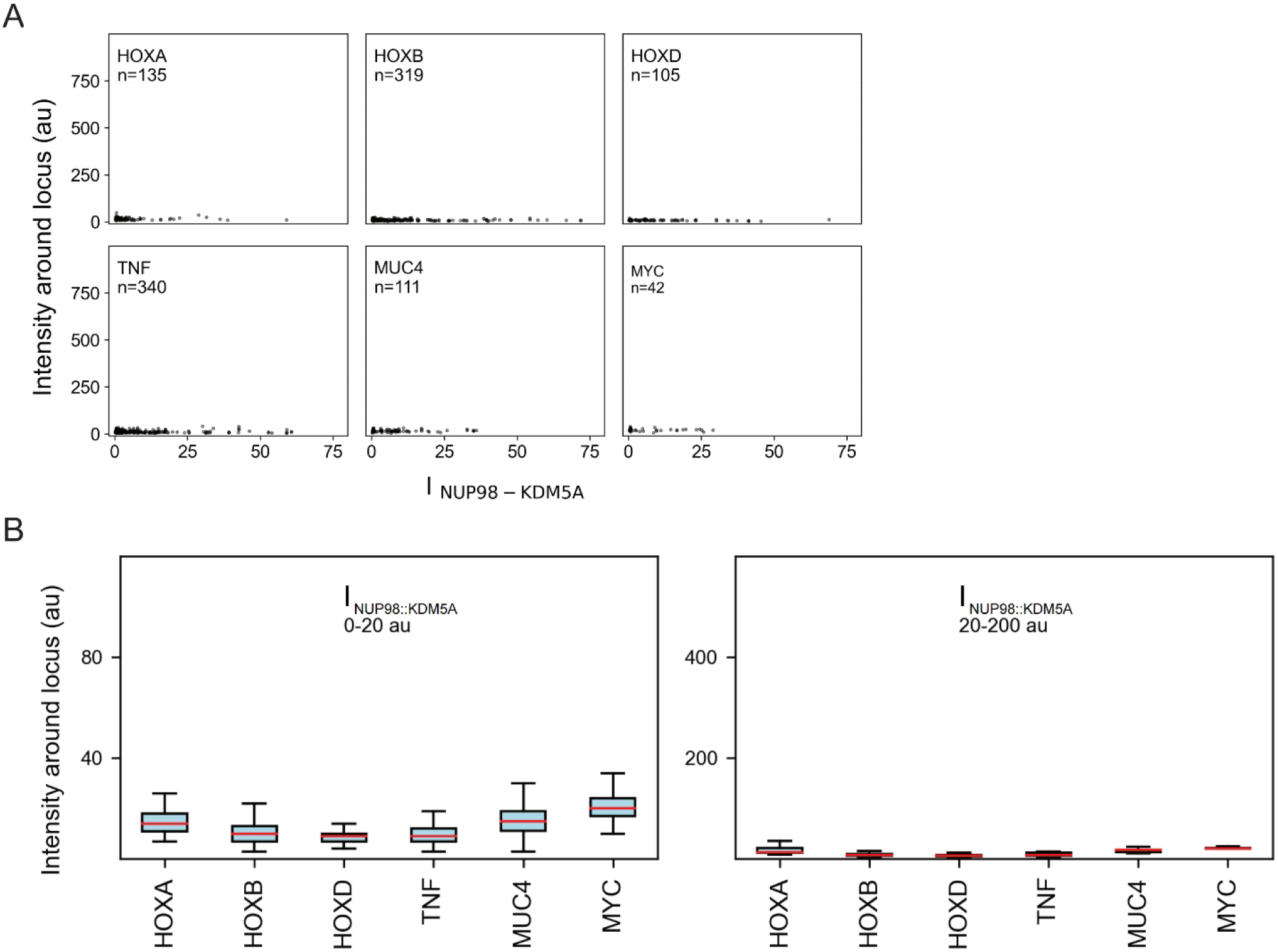
Negative control for the local signal analysis. **A.** Intensity around the FISH-labeled loci (HOXA, HOXB, HOXD, TNF, MUC4, and MYC) on the DAPI channel across a range of NUP98::KDM5A-FLAG expression levels in HEK293FT cells. **B.** Boxplots showing the DAPI channel intensity surrounding FISH-labeled loci (HOXA, HOXB, HOXD, TNF, MUC4, and MYC) across binned NUP98::KDM5A-FLAG expression levels in HEK293FT cells.

**Figure 4 Figure Supplement 1.**
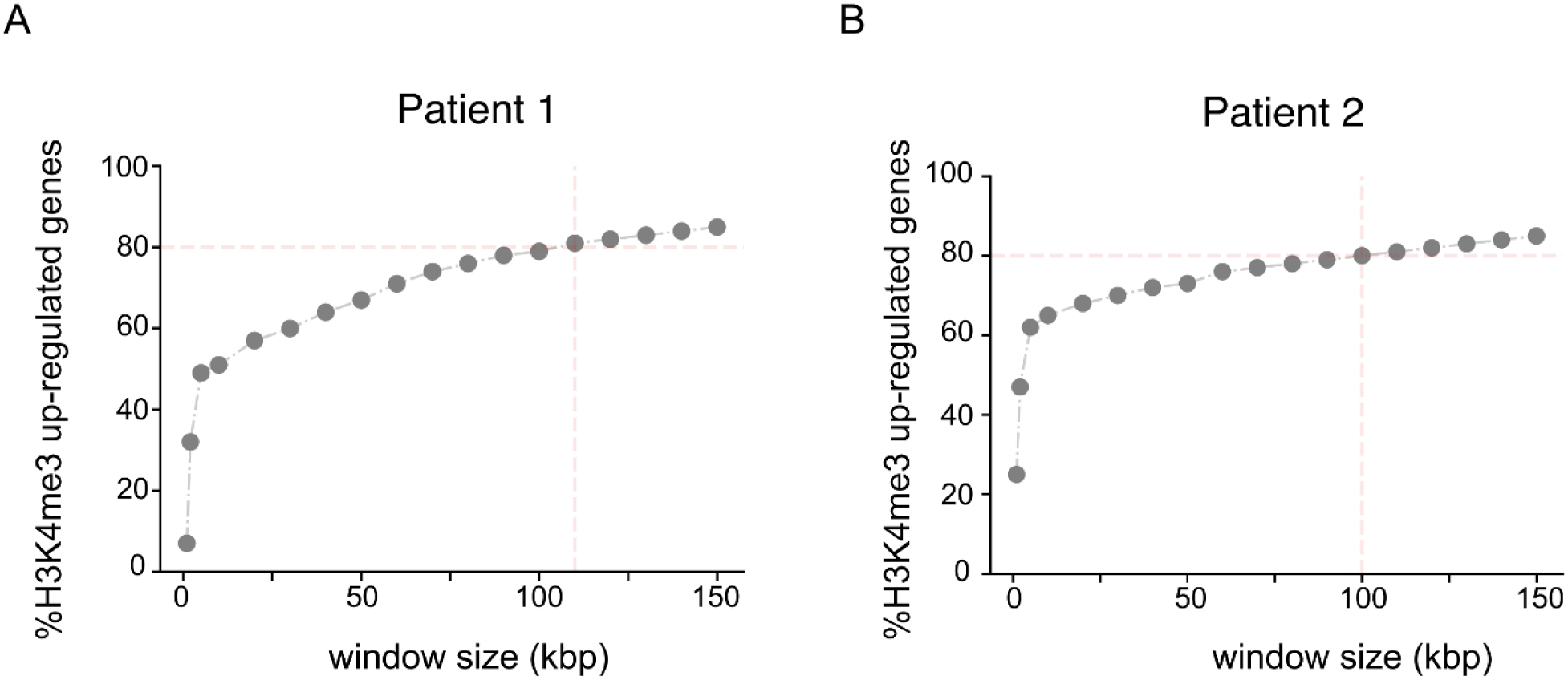
Selection of window size for H3K4me3 quantification. Window size selection for the quantification of gene-specific H3K4me3 intensity.

**Supplementary Table 1.**

| Construct | Plasmid | Used in Figures |
| --- | --- | --- |
| NUP98(1-513)-KDM5A(1486-1690)-mEGFP | MTK0-pGK | Figure 1, Figure 2 |
| mEGFP-NUP98(1-513)-KDM5A(1486-1690) | MTK0-pGK | Figure 2 |
| NUP98(1-513)-KDM5A<sub>mut</sub>(1486-1690)-mEGFP | MTK0-pGK | Figure 1 |
| NUP98(1-513)-KDM5A(1486-1690)-mCherry | MTK0-pGK | Figure 1 |
| NUP98(1-513)-KDM5A(1486-1690)-FLAG | MTK0-pGK | Figure 3 |
| mEGFP-KDM5A(full length) | MTK0-pGK | Figure 1 |
| mEGFP-KDM5A<sub>mut</sub>(full length) | MTK0-pGK | Figure 1 |
| mEGFP-KDM5A(1486-1690) | MTK0-pGK | Figure 1 |

**Supplementary Table 2.**

| Construct | Plasmid | Used in figures |
| --- | --- | --- |
| 6xHis-NUP98 (1-513)-KDM5A (1486-1690) | pET-28a | Figure 2 |
| 6xHis-NUP98 (1-513)-KDM5A (1486-1690) W1625A | pET-28a | Figure 2 |
| 6xHis-MBP-PHD3 (1601-1662) | pET-28a | Figure 2 |
| 6xHis-NUP98 (1-513)-KDM5A (1601-1662) | pET-28a | Figure 2 |
| 6xHis-NUP98 (1-513)-KDM5A (1486-1600) | pET-28a | Figure 2 |
| 6xHis-NUP98 (1-513) | pET-28a | Figure 2 |

